# A multiplexed striatal architecture for generalized spatial goal progress

**DOI:** 10.64898/2026.04.08.716650

**Authors:** Norihiro Takakuwa, Zahra Golipour, Hiroshi T Ito

**Affiliations:** Department of Fundamental Neuroscience, University of Lausanne, Lausanne, Switzerland; Max Planck Institute for Brain Research, Frankfurt am Main, Germany

## Abstract

Flexible goal-directed behavior requires a generalized internal metric that tracks progress toward goals across locations, routes, and behavioral contexts. Although hippocampal–entorhinal cognitive maps encode detailed spatial representations, whether the brain constructs an abstract distance-to-goal state that generalizes across arbitrary start–goal combinations—a prerequisite for reinforcement learning over space—remains unknown. Here we show that the nucleus accumbens (NAc) computes such generalized spatial states. Using large-scale population recordings and causal perturbations in rats navigating toward dynamically changing goals, we find that NAc activity encodes a scale-invariant distance-to-goal signal, normalized by total journey length, across maze geometries and task rules. A decoder trained on this signal transfers to novel journeys, tracking physical path length rather than remaining time. This signal emerges selectively during memory-guided but not cue-guided navigation, dissociating spatial distance-to-goal coding from purely motivational or value representations. It persists during hippocampal and medial entorhinal cortex silencing but depends critically on dopaminergic input from the ventral tegmental area, which is necessary for accurate goal targeting. Beyond the current goal, NAc populations encode distances to previous goals within orthogonal subspaces, enabling parallel evaluation of counterfactual spatial states without interference. Consistent with this multiplexed architecture, reward omission reinstates search behavior toward former goals, whereas suppressing dopamine release in the NAc abolishes this memory-guided reinstatement. These findings establish the NAc as a substrate for generalized, multiplexed spatial state abstraction, revealing a striatal computation that bridges cognitive mapping and reinforcement learning to enable flexible goal-directed behavior in dynamic environments.

---

Flexible navigation requires animals to continuously track their spatial progress toward goals and adapt behavior as circumstances change. Across species, animals dynamically re-evaluate navigational strategies during ongoing journeys^1–4^, and distance-to-goal is a central variable supporting these adjustments. Indeed, studies spanning insects to mammals have highlighted distance estimation as a fundamental component of navigation^5–7^. However, in natural environments, goals change, paths vary, and spatial layouts rearrange. How the brain represents progress toward goals in a way that generalizes across such changing spatial contexts remains poorly understood.

In mammals, the discovery of allocentric spatial representations in the hippocampus and entorhinal cortex^8,9^ has motivated the idea that distance information is derived from internal cognitive maps^10,11^. Supporting this view, subsets of hippocampal neurons have been reported to correlate with distance to a fixed target, most prominently in Egyptian bats^12^, with similar signals observed in rodents^13–15^. A key challenge for such representations, however, lies in their generalization across locations. Changes in start or goal location are accompanied by remapping or modulation of hippocampal–entorhinal spatial codes^16–20^, leaving unresolved the question of how the brain could compute distance between arbitrary pairs of locations in a coordinate-invariant manner.

Furthermore, navigation unfolds as an extended sequence of decisions, and tracking distance only to the currently selected goal is often insufficient. Animals must not only approach newly defined targets but also avoid, bypass, or revisit prior locations^21–23^. Although neural correlates of current goals have been described^24–27^, these signals typically update entirely in accordance with the animal’s ongoing decisions, raising the question of whether the brain can maintain latent representations of previously visited locations while pursuing a new goal. Importantly, such retrospective information is not merely mnemonic. It requires the active evaluation of counterfactual states—unchosen or previously relevant goals that could become behaviorally relevant again. Whether ongoing and counterfactual information can be maintained in parallel within the same neural population represents a central challenge for understanding flexible goal-directed behavior.

The dopamine–striatum system has classically been studied for its roles in reward prediction, value learning, and action selection^28–31^, but accumulating evidence indicates that it also contributes to spatial navigation. Activity of dopaminergic neurons in the ventral tegmental area, as well as dopamine release in the nucleus accumbens (NAc), increases as animals approach distant rewards^32–34^, and pharmacological disruption of dopamine signaling in the NAc impairs navigational performance^35,36^. These findings have led to the prevailing view that dopamine signals provide motivational or value-related information that supports goal-directed navigation. Dopamine ramping signals observed during navigation are often interpreted within temporal-difference (TD) reinforcement learning frameworks^37^, in which prediction errors are usually defined over elapsed time to reward^33,38–40^. A central strength of this framework is its capacity to generalize across diverse time-based tasks. Navigation, however, requires learning and inference over spatial state variables rather than time per se. Moreover, although reinforcement-learning frameworks can in principle incorporate information about previously rewarded locations through latent state representations^41^, whether and how such spatial and mnemonic variables are encoded within neural states remains unknown. Despite extensive theoretical work, the neural substrate for reinforcement learning over space—capable of computing distance between arbitrary pairs of locations and maintaining latent representations of prior goals—has not yet been identified, leaving a critical gap between cognitive map theories and spatial learning mechanisms in the brain.

Here we address this gap by developing a navigation task that dissociates spatial state from instantaneous position, trajectory, and movement direction while allowing systematic manipulation of goal identity. Using large-scale Neuropixels^42,43^ recordings, fiber photometry to monitor dopamine concentration, and pharmacogenetic and optogenetic perturbations, we test whether the NAc serves as a substrate for computing generalized spatial states defined by physical distance to the goal, as predicted by reinforcement-learning frameworks operating over space. By varying goal locations and running directions, we examine the generality and scalability of these representations. Critically, the task also enables examination of how previously visited goal locations are represented because animals must traverse them en route to a new target, providing direct access to neural representations of unchosen counterfactual options that are difficult to isolate in conventional discrete cue-choice tasks. Together, this approach reveals a multiplexed spatial state architecture in the NAc that integrates current and prior goal information within low-dimensional ensemble states, establishing a circuit basis for flexible navigation in dynamic environments.

## Shared goal-progress dynamics in NAc

To examine neuronal population activity in the nucleus accumbens (NAc) during goal-directed spatial navigation, rats were trained on a linear-maze task in which they alternated between paired goal wells (Fig. 1a). One goal well was fixed at the end of the track, whereas the second goal systematically varied across trial blocks. Each block consisted of 10–20 trials, with each trial comprising an outbound and an inbound journey. At the start of a new block, a goal location was introduced by delivering a water reward while LEDs beneath all wells were illuminated, signaling the change in goal location. In subsequent trials, rats were required to lick at the correct well for a fixed duration (1–1.5 s) to obtain a reward. All wells were identical in shape, and experiments were conducted under minimal lighting conditions to discourage reliance on local sensory cues for goal identification. This design ensured that rats engaged in goal-directed navigation from the onset of each journey rather than random foraging, as reflected by smooth uninterrupted running speed profiles without mid-journey slowing (Fig. 1a).

**Figure 1.**
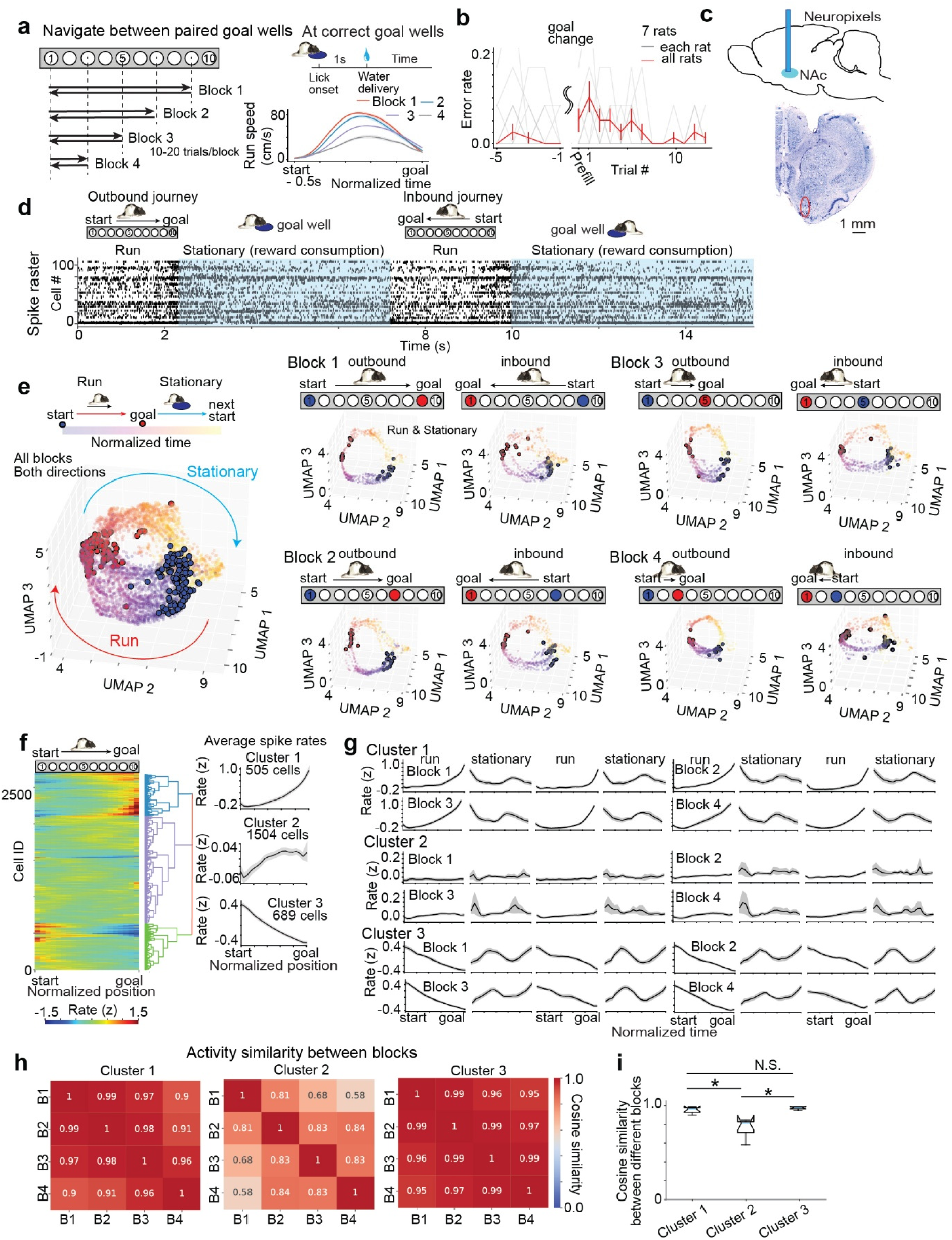
Goal-invariant evolution of NAc population dynamics during navigation. **a,** Left: Linear-maze task schematic. Rats alternated between goal wells for 10–20 trials per block; one goal shifted nearer each block. Reward required >1 s sustained licking. Right bottom: Mean running speed by normalized position (mean ± s.e.m.; 26 sessions, 7 rats). **b,** Error rate around goal changes for journeys toward the updated goal. Gray: individual animals; red: mean ± s.e.m. **c,** Sagittal schematic (top) and coronal section (bottom) showing Neuropixels probe placement in NAc. Red circle: probe tip. Scale bar, 1 mm. **d,** Spike raster of NAc population activity during a representative trial (outbound run, reward, inbound run, reward). Blue shading: stationary reward-consumption periods. **e,** UMAP embedding of NAc population activity. Left: All trials from a representative session, colored by journey progression (purple → yellow). Blue/red outlined points: motion onset/goal arrival. Right: Per-block outbound and inbound embeddings, showing preserved ring-like structure across configurations. **f,** Hierarchical clustering of NAc firing patterns between start and goal. Left: z-scored firing-rate matrix. Middle: dendrogram. Right: mean cluster profiles (26 sessions, 7 rats, n = 2,698 neurons). **g,** Cluster firing-rate profiles per block and trial for each task phase (mean ± s.e.m.). **h,** Cosine similarity of firing-rate profiles between blocks per cluster, showing block-wise consistency in Clusters 1 and 3 but not Cluster 2. **i,** Cosine similarity comparison across clusters. *p < 0.05, Wilcoxon rank-sum tests (Clusters 1 vs. 3: p = 0.200; 2 vs. 3: p = 0.004; 1 vs. 2: p = 0.004).

Although task performance transiently declined following changes in goal location, it recovered within one to two trials (Fig. 1b), consistent with efficient and flexible goal updating. To investigate how this rapid updating is reflected in and implemented by neural activity, we implanted Neuropixels probes in the nucleus accumbens (NAc) to enable simultaneous recordings from large populations of neurons (45–149 cells, 103.8 ± 4.8 per session; Fig. 1c and Extended Data Fig. 1). Recording tracks spanned the NAc, with most probes traversing the core region based on histological inspection. Core and shell neurons were not separated, and all analyses were conducted on the pooled population (Extended Data Fig. 1).

Despite substantial trial-to-trial variability in navigation length, direction, and time duration, we observed a consistent pattern of neural activity across trials and blocks (Fig. 1d). To confirm this observation, we applied uniform manifold approximation and projection (UMAP)^44^ to NAc population activity spanning both running and stationary reward-consumption phases. In this low-dimensional space, population activity evolved along ring-shaped trajectories, advancing from navigation onset toward goal arrival and returning toward the origin during the stationary phase (Fig. 1e). Persistent homology analysis applied to the UMAP embeddings confirmed that population activity formed a single connected structure in each block (one infinite H0 feature) and exhibited a significant one-dimensional topological signature (H1) relative to shuffle controls (Extended Data Fig. 2), consistent with the ring geometry. This topological signature was preserved across individual goal blocks and running directions, confirming shared population dynamics across navigation trials.

To link these population-level goal-directed dynamics to single-neuron activity, we classified the firing patterns of individual NAc neurons during navigation across trials and blocks using hierarchical clustering (Fig. 1f). The optimal number of clusters was determined to be three using the elbow method (see Methods). Examination of cluster-averaged activity profiles revealed two block-invariant patterns. Neurons in Cluster 1 showed a progressive increase in activity as animals approached the goal, whereas neurons in Cluster 3 exhibited a progressive decrease, and both patterns were highly consistent across blocks. In contrast, neurons in Cluster 2 also showed goal-approach-related increases in activity, but these patterns varied across blocks (Fig. 1g).

To quantify block-wise consistency, we computed the cosine similarity of population activity patterns between blocks for each cluster (Fig. 1h). For each cluster, the population activity was averaged across neurons within the cluster for each block, separately for the inbound run, rest, outbound run, and subsequent rest epochs. Cosine similarity was then computed between pairs of these block-wise population-averaged activities. Clusters 1 and 3 exhibited highly consistent activity across blocks (mean cosine similarity = 0.973 and 0.952, respectively), whereas Cluster 2 showed substantially lower between-block similarity (0.763). Direct cluster-wise comparisons revealed that Cluster 2 exhibited significantly lower similarity than both Cluster 1 and Cluster 3 (Cluster 1 vs. 2, p = 0.0039; Cluster 2 vs. 3, p = 0.0039), while Clusters 1 and 3 did not differ significantly from each other (Clusters 1 and 3, p = 0.2002; Wilcoxon rank-sum test; Fig. 1i), indicating greater block-dependent variability in Cluster 2.

Together, these results demonstrate that block-invariant goal-directed population dynamics in the NAc are predominantly supported by neurons exhibiting monotonic ramp-up (Cluster 1) and ramp-down (Cluster 3) activity patterns. At the ensemble level, NAc ensemble activity evolves along a shared population trajectory during goal-directed navigation that is largely invariant to goal location and running direction.

## Generalized distance-to-goal coding in NAc

We next asked which behavioral variables are captured by the shared goal-directed dynamics in the NAc. Based on previous work on the dopamine–striatal system, one possibility is that this activity reflects state-value signals that increase as animals approach a reward^32,38^. If NAc neurons contribute to such state representations, their activity may explicitly encode proximity to the goal, either in temporal or spatial terms. To test this possibility, we examined whether NAc population activity represents goal proximity using a decoding approach.

Goal proximity can be parameterized in multiple ways, including elapsed time, traversed distance, or these quantities normalized by the journey’s total duration or length. We therefore systematically tested these four candidate proximity variables (Fig. 2a and Methods). Decoding was performed using linear support vector regression trained on data from two task blocks, including both running directions, and evaluated on the remaining two blocks with different goal locations. By training and testing on distinct blocks, this approach provided an explicit test of whether proximity representations generalized across start-goal combinations.

**Figure 2.**
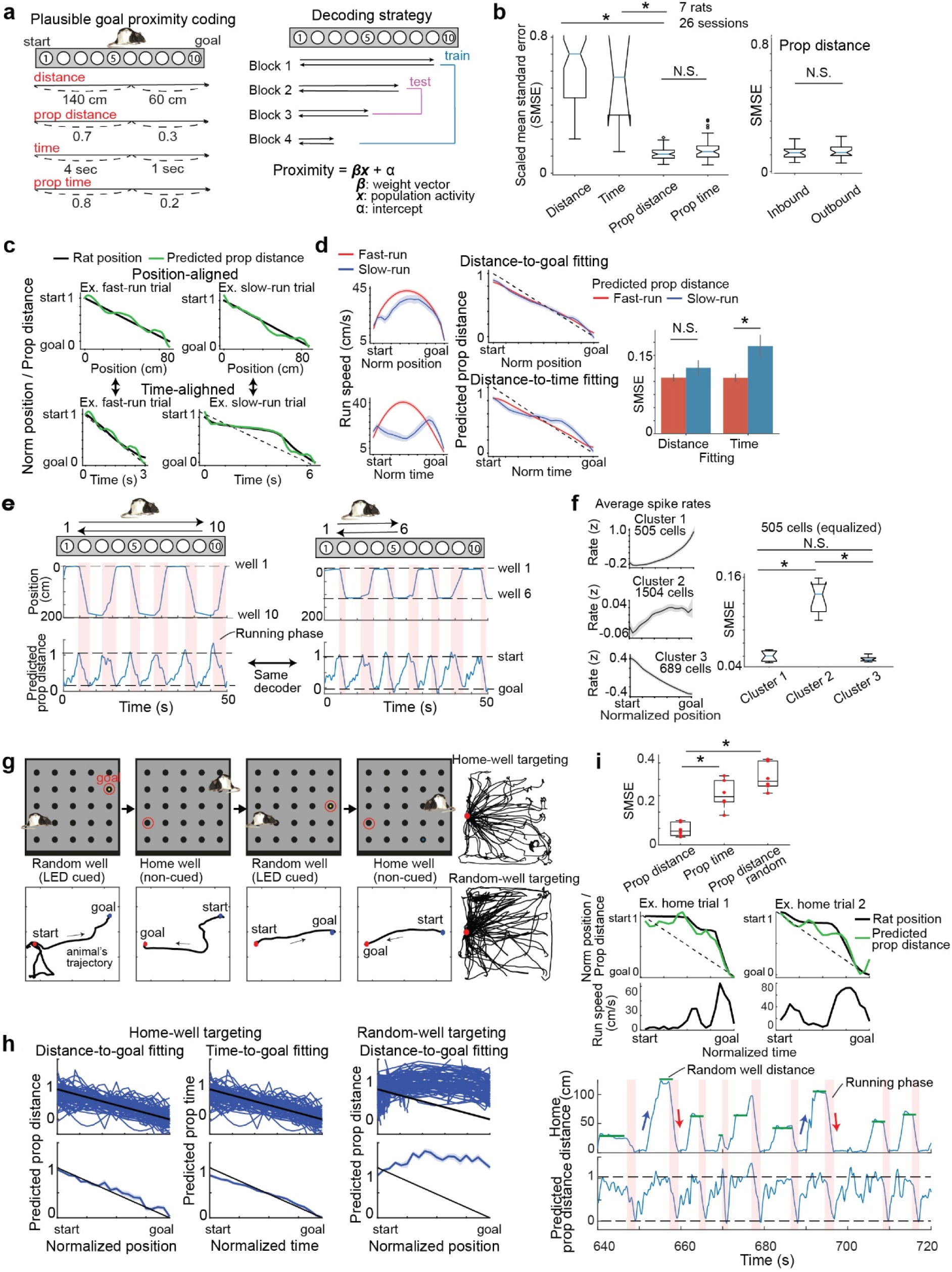
NAc activity tracks proportional distance to the goal in one- and two-dimensional environments. **a,** Left: Candidate progress variables: distance, time, proportional distance, and proportional time (normalized to total journey length or duration, respectively). Right: Linear decoders trained on outbound and inbound runs from two of four blocks and tested on the remaining two (all combinations). **b,** Left: Scaled mean squared error (SMSE) was significantly lower for proportional distance than distance or time (p < 0.001, Wilcoxon rank-sum test) but equivalent to proportional time (p = 0.964). Right: No significant difference between outbound and inbound proportional distance decoding (p = 0.560). **c,** Representative fast- and slow-run trials with proportional distance decoding fitted against distance or time. Dashed lines: proportional time. Note deviation from diagonal in the time-aligned slow-run plot. **d,** Averaged fast-run (red; 3,188 trials) and slow-run (blue; 106 trials) analysis. Left: Speed profiles against normalized distance (top) or time (bottom); mean ± s.e.m. Middle: Predicted proportional distance against normalized distance or time; dashed lines: identity. Right: SMSE comparison where decoding errors differed significantly for time (p = 0.007) but not distance (p = 0.233). **e,** NAc distance coding across consecutive trials. Top: animal position. Bottom: decoded proportional distance. Red shading: journey epochs; dashed lines: start/goal. Trials before (left) and after (right) goal change demonstrate generalized proximity coding across configurations. **f,** Left: Mean activity of the three clusters (Fig. 1f) along normalized position. Right: Clusters 1 and 3 showed significantly lower SMSE than Cluster 2 (p < 0.001); Clusters 1 vs. 3 did not differ (p = 0.386). **g,** Two-dimensional navigation task in a square maze (5 × 5 well grid). Rats alternated between random-well-targeting (LED-cued) and home-well-targeting (memory-guided) phases. Top panels: example trial schematics (rat position = start; red circle = target). Bottom panels: corresponding trajectories. Right: Example session trajectories for home-well (top) and random-well (bottom) trials; home well in red. **h,** Goal proximity decoding from NAc activity. Decoder trained on home-well trials. Top: individual trial performance against normalized position or time (black diagonal = identity). Bottom: mean ± s.e.m. Left: proportional path distance (home-well). Middle: proportional time (home-well). Right: proportional path distance (random-well). **i,** Top: Proportional distance during home-well trials was decoded more accurately than proportional time or proportional distance during random-well trials (SMSE: 0.082 ± 0.015, 0.163 ± 0.041, 0.308 ± 0.057, respectively; *p < 0.05, Wilcoxon signed-rank). Middle: Two example home-well trials showing predicted proportional distance (green) against normalized time, with deviations from proportional time (dashed); lower panels: corresponding speed. Bottom: Continuous NAc distance coding: animal distance to home well (top) and decoded proportional distance (bottom). Blue arrows: random-well phases; red arrows: home-well phases; green: random-well locations. Data from 2 rats, 6 sessions, 434 neurons.

Decoder performance was quantified using the scaled mean squared error (SMSE) between predicted and true proximity values. Among the tested variables, proportional distance and proportional time yielded the lowest SMSEs, indicating that NAc activity represents goal proximity using a normalized state variable rather than absolute time or distance, supporting generalization across goal locations and running directions (Fig. 2a right; Fig. 2b) with consistent results across individual rats (Extended Data Fig. 2).

Although decoder performance did not differ significantly between proportional distance and proportional time, we reasoned that this similarity could arise from stereotyped navigation behavior in the task, which tightly couples elapsed time and traveled distance. To dissociate these variables, we compared two groups of journeys in which animals exhibited significantly different running-speed profiles, characterized by pronounced slowing in the middle of the trajectory (Fig. 2c). Importantly, these trials were not associated with behavioral errors or task disengagement. Under these conditions, the proportional-distance decoder accurately tracked normalized distance to the goal regardless of changes in running speed, whereas its ability to track normalized time deteriorated markedly, particularly on trials with mid-journey slowing (Fig. 2d). We quantified this effect using a two-way repeated-measures ANOVA on SMSE with factors of speed (fast vs. slow) and fitted axis (distance vs. time), which revealed a significant speed × axis interaction (p = 0.024). Post hoc comparisons showed that slow-run trials increased decoding error along the time axis (p = 0.007, Wilcoxon rank-sum test), but not along the distance axis (p = 0.233). These results indicate that proportional distance-to-goal decoding by NAc neurons cannot be explained by time as a confounding factor, but instead reliably tracks the animal’s spatial state irrespective of movement speed, supporting proportional distance as a state variable encoded in the NAc.

Given that NAc population activity encodes proportional goal distance, it remains unclear how this signal is maintained across repeated navigation trials that include stationary reward-consumption phases. We therefore examined how NAc neurons represent distance over extended periods encompassing both running and stationary phases (Fig. 2e). We found that, while the distance decoder reliably tracked goal proximity during individual journeys, the decoded signal reset to the farthest distance before the onset of the subsequent journey. Moreover, applying the same distance decoder across different goal blocks revealed that proximity could be accurately decoded irrespective of goal identity (Fig. 2e, right). These results indicate that goal-distance representations in the NAc are dynamically reset between trials and generalize across goals, consistent with the ring-shaped low-dimensional manifold dynamics observed in Fig. 1.

This observation motivated us to examine the contributions of individual neurons to the population-level dynamics. We assessed the contributions of the three neuronal clusters identified in Fig. 1f and found that distance-to-goal decoding was predominantly supported by Clusters 1 and 3, which exhibited goal-directed ramp-up or ramp-down firing as animals approached the goal, respectively (Fig. 2f). In contrast, Cluster 2 showed poor decoding performance, indicating that goal-distance representations in the NAc are primarily carried by neurons whose firing rates change monotonically with distance to the goal.

Finally, given previous reports of goal-distance coding in the hippocampus to a fixed destination^12,13^, we asked whether the hippocampus also exhibits goal-invariant distance representations similar to those observed in the NAc, using the same decoding approach. We recorded neuronal activity from animals implanted with a tetrode microdrive in hippocampal CA1 while they performed a goal-directed navigation task on the linear maze and applied an identical distance-decoding analysis across different start-goal combinations (Extended Data Fig. 2). In contrast to the high decoding accuracy observed in NAc populations, decoding errors in CA1 were substantially larger under matched conditions, indicating that hippocampal distance representations are not generalized across start-goal combinations. These results suggest that goal-invariant proportional distance coding is a distinctive feature of NAc ensemble activity.

## Two-dimensional goal distance coding in NAc

The robust distance coding observed in the NAc during linear-maze navigation motivated us to examine whether and how this representation generalizes to a two-dimensional environment (Fig. 2g). To address this question, we employed a modified version of the task introduced by Pfeiffer and Foster (2013)^27^, in which animals alternated between cue-guided navigation to a randomly-assigned well and memory-guided targeting to a fixed home well (see Methods). The maze contained 25 wells, with the home well fixed throughout the session. During cue-guided navigation, animals navigated to a randomly assigned target well indicated by an LED cue. After reward consumption at the cued location, animals returned to the home well in the absence of explicit sensory cues, requiring navigation based on memory of the home location. This task therefore differed from the linear-maze paradigm in both spatial layout and behavioral rule structure.

We applied the same decoding framework to NAc population activity during this task. For each home-directed journey, we computed the proportional distance from the starting well—randomly assigned on each trial—to the fixed home well and used this variable to train the decoder. When applied to held-out trials, the decoder accurately predicted proportional goal distance during individual home-directed trajectories, despite substantial trial-by-trial variation in starting locations (Fig. 2g,h; Extended Data Fig. 2). Importantly, in the two-dimensional maze, Euclidean distance and path distance diverge depending on the animal’s chosen trajectory. The decoder more closely tracked path distance than Euclidean distance (Extended Data Fig. 2), consistent with an internally generated estimate of spatial progress derived from self-motion signals rather than simple geometric proximity. In contrast, the same decoder failed to predict proportional distance during cue-guided navigation toward randomly-assigned cued wells (proportional distance home vs. random: p = 0.031; Wilcoxon signed-rank test), confirming that NAc distance coding does not simply reflect reward approach, habitual activity, or motivational drive, but instead arises from active computation during ongoing navigation and is selectively expressed during memory-guided goal-directed behavior toward a remembered goal (Fig. 2h,i).

Consistent with our linear maze results, decoding of proportional time performed significantly worse than decoding of proportional distance (Fig. 2h,i; proportional distance vs. proportional time: p = 0.031; Wilcoxon signed-rank test). Moreover, neither head direction nor goal direction could be reliably decoded from NAc activity (Extended Data Fig. 2). Together, these results extend our linear-maze findings to a two-dimensional spatial layout with a distinct task rule, demonstrating that NAc spatial progress coding generalizes across environments and task structures. This coding is selectively expressed during memory-guided navigation and is independent of elapsed time or directional signals.

## NAc distance coding is independent of HPC and MEC

The finding that NAc activity accurately tracks distance to the goal raises the possibility that this coding relies on cognitive map representations described in the hippocampus (HPC) and the medial entorhinal cortex (MEC)^8,9^. To test this hypothesis, we performed pharmacogenetic inhibition of neurons along the dorsal–ventral axis of the HPC as well as in the MEC by expressing the inhibitory DREADD hM4Di^45^ (Fig. 3a; Extended Data Fig. 3). Neuronal activity was suppressed by subcutaneous injection of the DREADD agonist deschloroclozapine (DCZ)^46^ and the effectiveness and temporal profile of inhibition were verified following DCZ administration (Extended Data Fig. 4). The functional impact of silencing was further validated behaviorally using an object location memory task (Extended Data Fig. 4), which depends on intact hippocampal function^47,48^. Whereas control animals preferentially explored the object displaced from its original location after a 1-hour delay, animals receiving DCZ injections failed to show this preference, indicating impaired object location memory consistent with effective hippocampal silencing.

**Figure 3.**
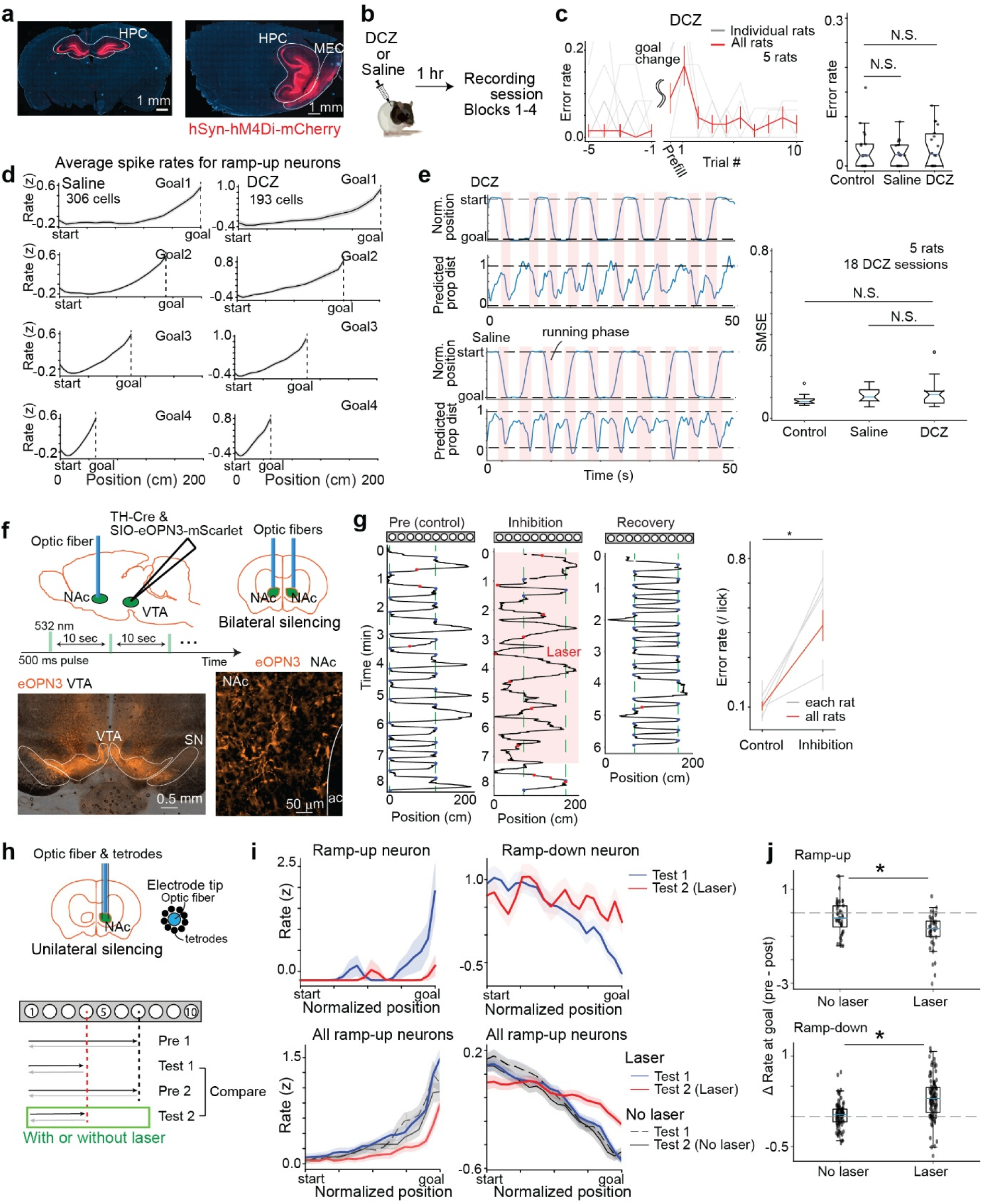
NAc distance coding requires dopaminergic input from the VTA but not hippocampal or medial entorhinal neuronal activity. **a,** Coronal and sagittal brain sections showing hM4Di expression in the hippocampus (HPC) and medial entorhinal cortex (MEC). Scale bar, 1 mm. **b,** Experimental design: systemic DCZ or saline administration 1 h before behavioral tasks. **c,** Left: Trial-by-trial error rates around goal changes for DCZ sessions. Right: Mean error rates across control, saline, and DCZ conditions showed no significant differences (control vs DCZ, p = 0.693; control vs saline, p = 0.703; DCZ vs saline, p = 0.754, Wilcoxon rank-sum; 5 rats; control: 20, saline: 17, DCZ: 18 sessions). **d,** Average firing rates of NAc neurons exhibiting goal-directed ramp-up activity across four task blocks with different goal locations, shown separately for saline and DCZ sessions. **e,** Left: Proportional distance coding dynamics across consecutive epochs for saline and DCZ sessions. Right: SMSE comparison revealed no significant differences between conditions (control vs DCZ, p = 0.090; DCZ vs. saline, p = 0.869; control vs saline, p = 0.054; Wilcoxon rank-sum test). **f,** Optogenetic inhibition design. AAV-eOPN3 was injected bilaterally into VTA under the control of tyrosine hydroxylase promoter with optic fibers in NAc. Histology shows eOPN3 expression in VTA and axonal projections in NAc. Scale bars, 0.5 mm (VTA), 50 µm (NAc). **g,** Bilateral dopaminergic inhibition effects. Left: Example trajectories during control, inhibition, and recovery; blue/red dots: correct/error licks. Right: Error rates increased significantly during inhibition (control: 10.4 ± 1.8%; inhibition: 42.9 ± 6.2%; p < 0.001; 16 sessions, 4 rats). **h,** Unilateral inhibition design with opto-tetrode microdrive. Four-block task controlled for goal-change effects: first goal change (Pre 1 → Test 1) served as control; second (Pre 2 → Test 2) compared laser-on vs. laser-off conditions. **i,** Firing rates of representative neurons (top) and population-averaged normalized rates (bottom) for ramp-up (left) and ramp-down (right) neurons by normalized position. Black: no-laser (dashed, Test 1; solid, Test 2); blue/red: laser (Test 1/Test 2). **j,** Firing-rate changes near the goal (last 2 of 20 position bins). Laser disrupted both ramp-up (p = 0.003; control: 40 cells, 2 rats, 8 sessions; inhibition: 36 cells, 2 rats, 10 sessions) and ramp-down firing (p < 0.001; control: 80 cells; inhibition: 96 cells; Wilcoxon rank-sum).

In the same animals, we implanted Neuropixels probes in the NAc to examine the impact of HPC and MEC inhibition on distance-to-goal coding. We systematically compared non-injected control, saline, and DCZ sessions within each animal. At the behavioral level, we did not detect differences in running speed or task performance across conditions (Fig. 3c; control vs DCZ, p = 0.693; control vs saline, p = 0.703; DCZ vs saline, p = 0.754; Wilcoxon rank-sum test; Extended Data Fig. 4). At the neural level, goal-directed ramp-up firing patterns in the NAc were preserved in both saline and DCZ sessions (Fig. 3d). Consistent with this observation, decoding performance of proportional goal distance did not differ between control, saline, and DCZ conditions (Fig. 3e; control vs DCZ, p = 0.090; DCZ vs saline, p = 0.869; control vs saline, p = 0.054; Wilcoxon rank-sum test). Together with the weak distance-to-goal coding observed in hippocampal neurons (Extended Data Fig. 2), these results indicate that NAc distance representations do not require intact HPC or MEC population activity during navigation, suggesting that NAc distance coding is not simply inherited from online hippocampal map representations.

## VTA dopamine is required for NAc distance coding

Dopaminergic neurons in the ventral tegmental area (VTA) provide a major input to the NAc^49,50^. We therefore next examined whether this dopaminergic input is required for distance-to-goal coding in the NAc. To selectively suppress inputs from VTA dopamine neurons, we injected an adeno-associated virus (AAV) encoding the inhibitory opsin eOPN3^51^, optimized for axon-terminal silencing, into the VTA. Dopamine neuron–specific expression was achieved using Cre recombinase driven by the tyrosine hydroxylase promoter (Fig. 3f). This approach yielded largely selective Cre-dependent opsin expression in tyrosine-hydroxylase-positive cells (87% of opsin-expressing cells were tyrosine-hydroxylase-positive; see Methods). Injections were performed bilaterally, and optical fibers were implanted bilaterally in the NAc to inhibit dopaminergic axon terminals originating from the VTA (Fig. 3f; Extended Data Fig. 1).

After animals were well trained on the task, they performed under conditions of bilateral inhibition of dopaminergic input to the NAc. This manipulation significantly impaired task performance, with animals frequently passing over the correct goal wells (Fig. 3g). Accordingly, the probability of choosing incorrect wells was markedly increased (control: 10.4 ± 1.8%, inhibition: 42.9 ± 6.2%; p < 0.001; Wilcoxon rank-sum test), indicating that dopaminergic input to the NAc is required for accurate goal targeting. This conclusion was further supported by direct silencing of VTA dopamine neurons, which produced comparable behavioral impairments (Extended Data Fig. 5). To determine whether these deficits specifically reflected impaired goal updating, we inhibited dopaminergic signaling after animals had already achieved stable goal-choice performance for a given goal pair. Under these conditions, silencing continued to impair performance, indicating that the observed deficits cannot be attributed solely to impaired learning of new goals but reflect a more general disruption of goal-directed navigation toward remembered goals (Extended Data Fig. 5).

To assess the contribution of VTA dopaminergic input to NAc distance coding independently of behavioral impairments, we sought experimental conditions under which dopaminergic silencing minimally affected task performance. We found that unilateral inhibition of VTA dopamine input to the NAc did not significantly disrupt navigation behavior (Extended Data Fig. 5), allowing us to examine neural consequences under matched behavioral conditions (Fig. 3h). We implanted a tetrode drive combined with an optic fiber in the NAc, enabling simultaneous single-unit recordings and optogenetic manipulation. Neurons exhibiting goal-directed ramp-up or ramp-down firing were first identified. To avoid confounds related to goal learning, neural activity was compared across two blocks with identical sequences of goal configurations (Fig. 3h).

Under these conditions, goal-directed modulation of NAc neurons was markedly disrupted at both the single-neuron and population levels (Fig. 3i). During dopaminergic silencing, ramp-up neurons failed to increase their firing rates and ramp-down neurons failed to decrease their firing as animals approached the goal (Fig. 3j). Together, these results demonstrate that while NAc distance-to-goal coding does not depend on hippocampal or medial entorhinal inputs, it critically requires dopaminergic input from the VTA.

## Multiplexed coding of current and previous goals

While NAc neurons accurately track distance to the currently targeted destination following goal changes, navigation often requires maintaining information about previously rewarded locations even after they are no longer behaviorally targeted. We therefore asked whether NAc neurons encode information about prior goals by designing a task in which animals sequentially navigated from the same start well to a distal well (Block 1), then to a proximal well (Block 2), and subsequently returned to the original distal well (Block 3; Fig. 4a). Because Blocks 1 and 3 shared the identical target well, this design allowed us to isolate the influence of experience with the intervening proximal-well targeting on neural activity while navigating to the same target.

**Figure 4.**
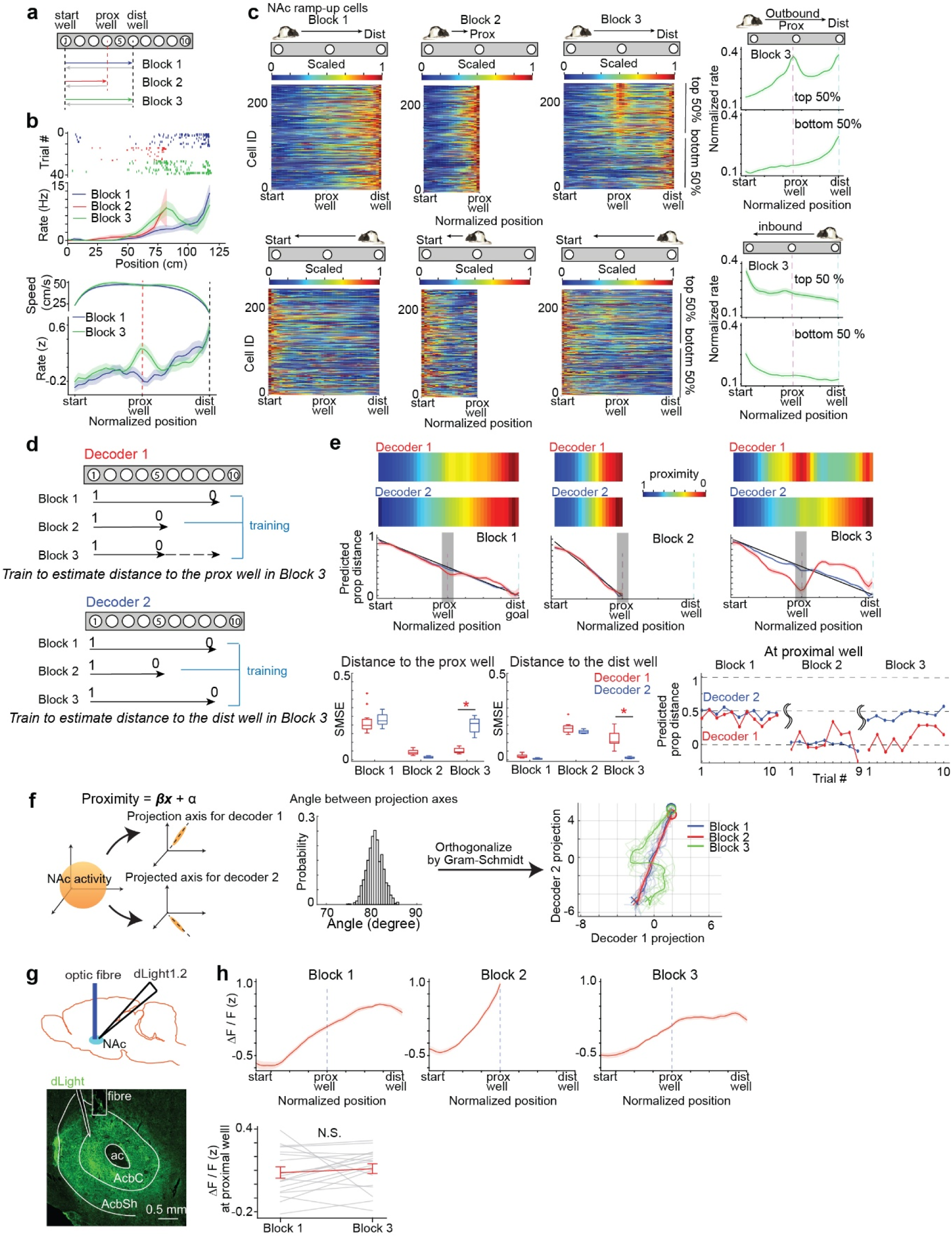
NAc neurons concurrently track spatial progress toward both current and previous goals along orthogonal subspaces. **a,** Task schematic: three blocks with sequential goal-well changes. One end-well was fixed; the paired goal alternated (Blocks 1 and 3: distal well; Block 2: proximal well). Data from 10 rats, 31 sessions, 2,209 neurons. **b,** Top: Representative neuron firing rates across blocks (outbound runs). Despite shared goals in Blocks 1 and 3, firing near the proximal well differed. Bottom: Population-averaged rates (top 25% neurons, 61 cells) under speed-matched trials (114 each) for Blocks 1 and 3. Red/black dashed lines: proximal/distal well. Firing rates differed without speed changes. **c,** Normalized firing rates of ramp-up neurons across blocks and directions, sorted by proximal-well activity in Block 3. Right: Mean rates for top and bottom 50% neurons. Previous-goal representations appeared during outbound but not inbound runs. **d,** Two-decoder strategy: both trained on Block 1 (distal well) and Block 2 (proximal well) proximity. In Block 3, Decoder 1 was trained on proximal-well (previous goal) and Decoder 2 on distal-well (current goal) proximity. Both used all ramp-up neurons (247 cells). **e,** Top: Decoded proportional goal proximity across blocks (outbound runs) as color-coded plots and mean ± s.e.m. Bottom left/middle: SMSEs relative to proximal (left) and distal (middle) wells. Both decoders tracked the same goal in Blocks 1–2; in Block 3, they diverged where Decoder 1 tracked the proximal well, Decoder 2 the distal well (p < 0.001, Wilcoxon signed-rank). Bottom right: Trial-wise predictions diverged in Block 3 and remained separated throughout. **f,** Left: Decoder projection axis schematic. Middle: Bootstrap-estimated angle between axes (80.77° ± 0.05°; n = 1,000). Right: Population activity projected onto Gram–Schmidt-orthogonalized decoder axes across blocks; thin lines: single trials; thick lines: trial averages. **g,** Fiber photometry design using dLight1.2 dopamine sensor in NAc, with histological verification. Scale bar, 0.5 mm. **h,** Top: Dopamine signal ramped toward the current goal in all blocks with no increase at the previous goal in Block 3. Bottom: Fluorescence around the proximal well (bins 18–22 of 40) showed no significant difference between Blocks 1 and 3 (p = 0.210, paired Wilcoxon signed-rank; 4 rats, 18 sessions).

At the single-neuron level, ramp-up neurons exhibited increasing activity as animals approached the target well in both Blocks 1 and 2. However, during Block 3 when the animal targeted the distal well after the intervening proximal-well block, a subset of neurons showed increased activity when animals passed the location of the former proximal goal, despite the fact that this location was no longer visited (Fig. 4b, top). This neural modulation was not accompanied by overt behavioral differences. We selected trials from Blocks 1 and 3 with closely matched running-speed profiles. Even under these behavior-matched conditions with nearly identical speed trajectories, neural activity diverged selectively at the location of the former proximal goal (Fig. 4b, bottom), indicating that this signal reflects memory of prior goal locations rather than differences in ongoing behavior.

At the population level, visualization of activity from all recorded ramp-up neurons across inbound and outbound trajectories revealed that, in Block 3, approximately half of the population exhibited elevated activity near the location of the previously rewarded proximal well (Fig. 4c). The remaining ramp-up neurons continued to show monotonic increases in activity toward the currently targeted distal well. Notably, neither group exhibited increased activity at the prior goal location during inbound trajectories in Block 3, indicating that past-goal encoding is direction-dependent. Moreover, this previous-goal coding activity cannot be attributed to egocentric goal-distance memory alone, as neurons consistently represented the same previous goal location despite differences in starting positions between blocks (Extended Data Fig. 6), suggesting that this goal memory is encoded in an allocentric coordinate frame. In contrast to the robust mnemonic signals observed in goal-directed ramp-up NAc neurons, representations of previous goals were markedly weaker among ramp-down neurons (Extended Data Fig. 7).

The presence of two distinct populations of NAc neurons, preferentially encoding either the current or the previous goals, suggests that the NAc independently represents distance to these two locations at the population level. To directly test this idea, we constructed two goal-distance decoders using support vector regression (SVR), each optimized to decode proportional distance to either the currently targeted goal or the previously rewarded goal (see Methods). The two decoders differed only in how Block 3 trials were labeled during training: distances were computed relative to either the proximal well for the previous goal decoder (Decoder 1) or the distal well for the current goal decoder (Decoder 2). Decoder performance was evaluated on trials withheld from training using leave-one-out cross-validation.

During the first two blocks, both decoders accurately tracked distance to the currently targeted goal—the distal well in Block 1 and the proximal well in Block 2—with no apparent difference in decoding performance. In contrast, in Block 3, the predictions of the two decoders diverged. The current-goal decoder (Decoder 2) tracked distance to the distal well that animals were actively targeting, whereas the prior-goal decoder (Decoder 1) continued to track distance to the proximal well that had been targeted in the preceding block (Fig. 4e).

Specifically, in Block 1, both decoders indicated that the proximal well was far from the goal, consistent with the distal well being behaviorally relevant. In Block 2, both decoders correctly identified the proximal well as the goal. However, in Block 3, the current-goal decoder (Decoder 2) reverted to signaling that the proximal well was still distant from the goal, whereas the prior-goal decoder (Decoder 1) continued to report the proximal well as close to the goal. Notably, this representation of the previous goal persisted for an extended period even after behavior had fully adapted to the reinstated distal goal (Fig. 4e).

This finding raised the question of how the same neural population activity can simultaneously track distance to two distinct locations. To address this, we analyzed the projection axes of the two decoders for the current and previous goals, and found that they were nearly orthogonal (mean angle: 80.77° ± 0.05°, bootstrap resampling with 1,000 iterations; Fig. 4f and Methods). After further orthogonalizing these axes using the Gram–Schmidt orthogonalization procedure, we projected NAc population activity onto the two-dimensional subspace spanned by the two decoding axes to visualize the evolving population dynamics during individual journeys (Fig. 4f). In this subspace, population dynamics were nearly identical in Blocks 1 and 2, evolving along a straight trajectory from start to goal despite differences in goal locations, consistent with a generalized goal-proximity coding in the NAc. In contrast, this structure changed markedly in Block 3, with trajectories exhibiting pronounced curvature during the middle of journeys while preserving the same start and end points. Importantly, depending on the projection axis, the same population dynamics in Block 3 could reflect proximity to either the currently targeted distal well or the previously rewarded proximal well, enabling parallel and independent representation of goal proximity for both current and prior goals.

We further asked whether the NAc’s goal memory extends to multiple previously rewarded wells. To address this, we designed a task in which animals sequentially navigated between three distinct goal-well pairs that shared a common well (Extended Data Fig. 8). In the final block, when animals targeted the most distal goal well, they necessarily traversed the locations of the two previously rewarded wells, allowing us to test whether NAc population activity could simultaneously represent proximity to multiple prior goal locations (Extended Data Fig. 8). We found that NAc neurons encoded not only the currently targeted goal, but that a subset preferentially represented previously visited goal wells. Notably, we also identified distinct neuronal subsets that selectively represented only one of the prior goal locations, indicating that goal memory in the NAc is not governed solely by time-dependent decay but is also location-specific.

This observation raised the possibility that the persistence of NAc goal memory depends on task engagement or behavioral relevance. To test this hypothesis, we used the same task structure as in Fig. 4a but introduced an inter-block rest period of approximately 1 min, during which animals were removed from the maze and placed on a pedestal (Extended Data Fig. 6). This brief interruption markedly reduced representations of previous goals. The proportion of NAc neurons exhibiting prior-goal coding was significantly decreased, and the two decoders trained to track proximity to the current and previous goals no longer showed clearly distinct performance, indicating a substantial weakening of prior-goal representations. Given that a 1-min rest is comparable to the duration of a single trial—well below the time scale over which the NAc can maintain goal representations in the absence of inter-block rest (Fig. 4e)—these results indicate that maintenance of prior-goal representations depends on continuous task engagement rather than passive time alone.

Finally, to determine whether this goal-memory signal is unique to NAc neuronal activity or already present in dopaminergic input from the ventral tegmental area (VTA), we measured dopamine concentrations in the NAc using fiber photometry by expressing the dopamine sensor dLight1.2 (Fig. 4g). Across all task blocks, dopamine levels increased as animals approached the currently targeted goal. However, during Block 3—following the intervening proximal-goal block—dopamine concentration did not show a significant increase at the location of the previous proximal goal (Fig. 4h), in contrast to the population activity observed in the NAc. We further corroborated this dissociation by recording spiking activity from VTA neurons (Extended Data Fig. 9). Although a subset of VTA neurons exhibited goal-directed ramp-up firing that enabled prediction of proportional distance to the currently targeted goal, analogous to NAc neurons, these VTA neurons showed markedly weaker representations of previous goals (Extended Data Fig. 10). Thus, while dopaminergic signals convey a generalized goal-proximity signal, the multiplexed memory of alternative goals emerges within NAc population dynamics rather than being directly inherited from dopaminergic input.

## Reinstatement of prior goal search requires NAc dopamine

Our findings suggest that the NAc serves as a pivotal substrate for encoding distance not only to the currently targeted goal but also to previously rewarded locations. Although prior studies have identified several brain regions whose activity correlates with the ongoing navigation target, the persistence of latent coding for previous goals, even after behavior has been fully updated to a new goal, emerges as a distinctive feature of NAc activity. Such mnemonic or latent-state representations are critical in natural environments, where expected rewards are not always available and behavioral goals must be flexibly updated among multiple plausible alternatives, including past choices. We thus asked whether disrupting NAc goal coding impairs memory-guided search for previously rewarded locations.

To test this idea, we mimicked a naturalistic situation in which reward was unexpectedly unavailable at a learned goal location (Fig. 5a). Animals were first trained to navigate between a pair of goal locations. After stable performance was achieved, one of the goal locations was changed, requiring animals to update their navigation to the new goal. Once performance stabilized under this new configuration, defined as five consecutive correct trials (Fig. 5b), reward delivery at the newly learned goal well was omitted despite correct licking behavior. This reward-omission period lasted for 2 minutes, during which animals were free to explore, search, and lick at any well in the maze. During reward omission, animals initially continued to visit and lick the correct goal well despite the absence of reward, but subsequently began to explore and sample other wells in the maze (Fig. 5c). At the end of the reward-omission period, a water reward was delivered at the correct goal location independent of licking, after which animals immediately recovered accurate task performance.

**Figure 5.**
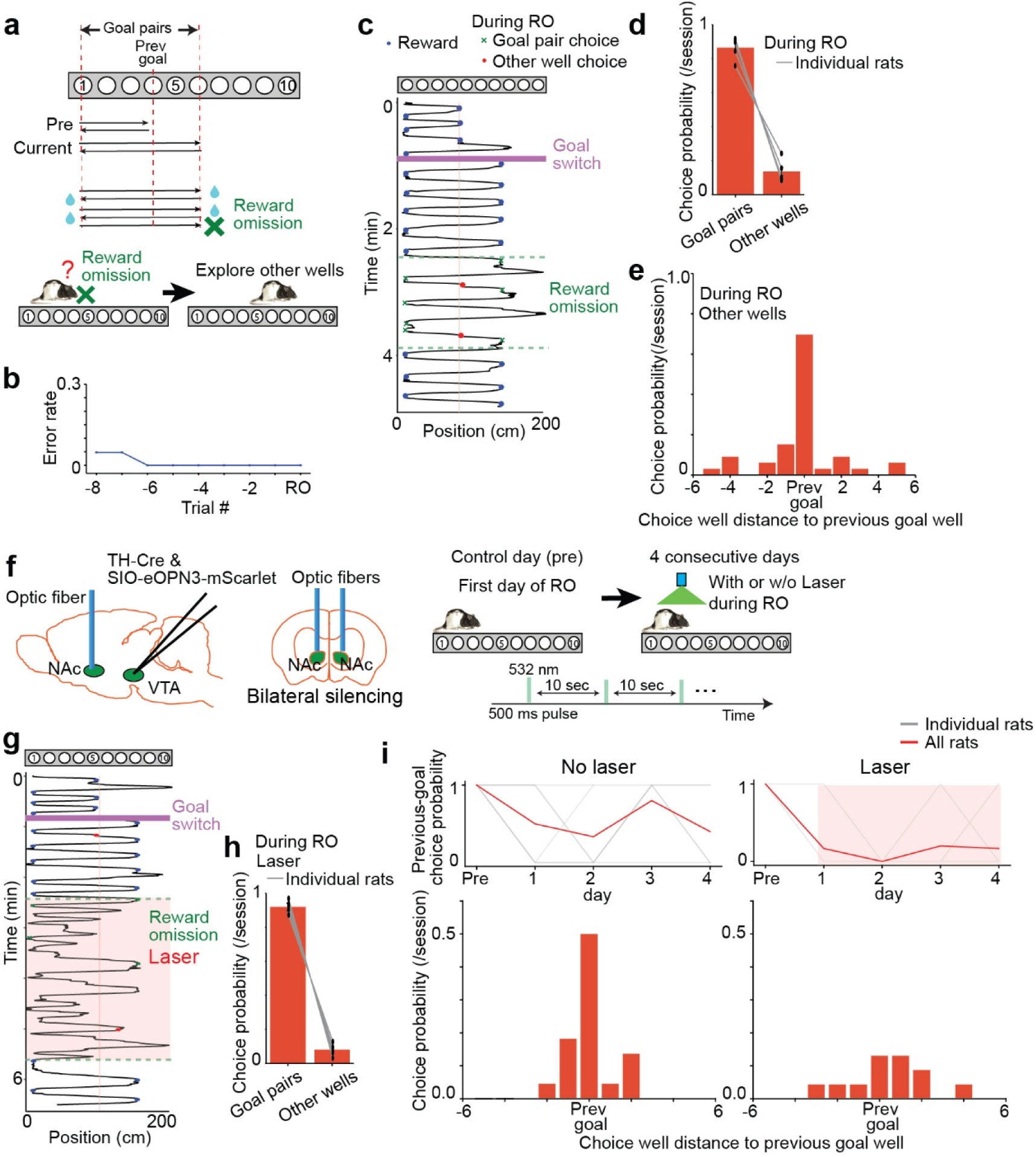
Inhibition of VTA-NAc dopaminergic input impaired the reinstatement of goal-directed search for previously learned targets. **a,** Reward-omission design. After learning a goal-well pair (≥5 consecutive correct trials), one goal was changed. After relearning, reward at the changed goal was omitted for ∼2 min, allowing assessment of exploratory choice behavior. Data from 6 rats, 33 sessions. **b,** Trial-wise error rates prior to reward omission, showing consecutive correct choices before omission. **c,** Animal behavior before and during omission. Red/blue dots: error/correct licks; green crosses: licks at correct goal during omission (no reward). Magenta bars: goal changes; green dashed lines: omission onset/offset. **d,** Well choice probability during omission: rats preferentially chose current goal-well pairs over other wells. **e,** Spatial distribution of non-goal choices during omission, grouped by distance from the previous goal. Choices were significantly biased toward the previous goal versus uniform random (p < 0.05, binomial test). **f,** Optogenetic inhibition design. Day 1 confirmed reliable previous-goal revisiting during omission. Over four subsequent days, omission sessions were conducted with or without laser. Pink shading: laser-on periods. **g,** Animal behavior during omission with laser application, as in **c**. **h,** Well choice probability during omission with laser: rats still preferentially chose current goal pairs despite reward absence. **i,** Top: Daily probability of visiting the previous goal during omission, with and without laser. Visits were binarized per session; gray lines: individual rats; red: mean over four test days. Bottom: Spatial distribution of choices during omission. Laser significantly reduced choices toward previously rewarded goals (p = 0.034, Wilcoxon rank-sum; no laser: 6 rats, 22 sessions; laser: 6 rats, 23 sessions), indicating impaired reinstatement of prior goal-directed behavior.

Analysis of animals’ behavior during the reward-omission period revealed that they continued to preferentially visit the current goal pair relative to other wells (Fig. 5d). However, when licking events outside the current goal pair were examined, animals showed a pronounced bias toward the well that had been rewarded in the previous block (Fig. 5e; p < 0.05; binomial test), indicating memory-guided search behavior.

Given this behavioral bias toward previously rewarded locations, we next asked whether memory-guided search depends on goal-memory signals in the NAc. Because goal-related coding in the NAc depends on dopaminergic input from the ventral tegmental area (VTA) (Fig. 3), we bilaterally suppressed dopaminergic input to the NAc specifically during the reward-omission period using the same optogenetic strategy as in Fig. 3 (Fig. 5f). To control for potential effects of repeated daily testing, reward-omission sessions were conducted in two separate five-day experiments with different rat groups, one without and one with dopaminergic inhibition (see Methods). On the first day, a baseline reward-omission session was performed to confirm preferential sampling of previously rewarded locations. On the subsequent four days, reward-omission sessions were conducted either with or without laser illumination.

Despite bilateral suppression of VTA–NAc dopaminergic input, animals continued to preferentially choose wells belonging to the current goal pair relative to other wells during the reward omission period (Fig. 5g,h), indicating that basic reward well recognition and well-licking behavior remained intact. Separate experiments confirmed that, although bilateral inhibition of VTA dopaminergic neurons after learning of a goal pair increased error rates, these errors largely reflected failures to alternate correctly between the goal-well pair, and animals still preferentially selected goal wells over non-goal wells (Extended Data Fig. 5). These findings suggest that VTA–NAc dopaminergic signaling is critical for flexible alternation once goal wells have been learned, but not for discrimination of learned goal locations from other locations. Notably, during the reward omission, these dopaminergic signals also contribute to memory-guided search for previously rewarded locations. We analyzed well choices made outside the current goal pair during the reward-omission phase. In control sessions, animals reliably revisited previously rewarded wells during reward omission at the current goals, as reflected by elevated choice frequency at the prior goal location (Fig. 5i). Under dopaminergic inhibition, however, this bias was abolished: despite preserved preferential selection of the current goal wells, choice frequency at the previous goal location was no longer elevated and was significantly reduced relative to control (p = 0.034; Wilcoxon rank-sum test; Fig. 5i). Collectively, these results indicate that dopaminergic input to the NAc is required for memory-guided selection of previously rewarded locations, supporting a role for the NAc in reinstating prior goals through multiplexed orthogonal coding of current and previous goal representations.

## Discussion

This study identifies a previously unrecognized goal-proximity computation in the NAc of the ventral striatum. Neurons in the NAc encode proportional distance-to-goal, normalized by total journey length, during navigation. This signal generalized across start–goal combinations and tracked physical approach by scaling with spatial displacement rather than elapsed time, providing a neural substrate for monitoring progress in spatial coordinates. The distance-to-goal coding was observed across distinct spatial layouts and task rules and depended critically on dopaminergic input from the VTA, yet persisted during silencing of hippocampal and medial entorhinal neurons, suggesting that the NAc can maintain progress signals independently of canonical hippocampal–entorhinal spatial representations. Beyond the current goal, NAc populations simultaneously compute distances to previously rewarded locations along orthogonal population axes, revealing a mnemonic component that preserves counterfactual options within the environment. Consistent with this multiplexed coding scheme, omission of expected reward biased search toward prior goal locations, whereas suppressing dopamine release in the NAc attenuated this memory-guided behavior. Together, these results identify the NAc as a neural substrate for estimating behavioral progress toward desired outcomes using a normalized distance metric, consistent with a spatial state variable that supports learning and decision-making over physical space.

Ramp-like activity in dopamine–striatal circuits has often been interpreted within temporal-difference (TD) learning frameworks as reflecting time-based prediction errors^28–30,33,38^. In contrast, the signals identified here tracked spatial proximity to internally represented goals in both one-and two-dimensional environments, remaining robust despite substantial variations in running speed, and were absent when targets were randomly assigned in cue-guided navigation. Together, these features argue against purely time-based or generalized motivational interpretations and instead indicate that NAc ramping reflects a spatially grounded computation of progress during internally driven navigation to remembered goals. More broadly, this interpretation is consistent with the idea that prediction error or value-related signals in dopamine–striatal circuits may be flexibly expressed over state variables defined in temporal or spatial domains.

Within reinforcement learning theories, dopaminergic signals are commonly interpreted as prediction errors over value estimates defined on abstract state spaces, which are usually temporally structured^28,29,37^. Our findings extend this framework by identifying a spatial state variable—distance-to-goal—that cannot be reduced to elapsed time, enabling reinforcement learning to operate over spatial state space. Normalizing distance by total journey length abstracts away from absolute spatial coordinates while preserving relational progress toward a goal, offering a solution to the challenge of generalization across changing start and goal locations within a given environment.

A distinctive feature of our navigation paradigm is that animals must traverse previously rewarded locations even after their target is updated—a condition not accessible in conventional cue-choice tasks. This design enables direct assessment of how neurons represent unchosen options. We found that NAc neurons encoded proximity not only to the currently selected target but also to previously visited goals, despite no overt behavioral expression of these alternatives, consistent with a mnemonic representation of latent goal states. Notably, distances to current and previous goals were encoded in orthogonal NAc population dimensions rather than integrated into a single composite state. Such orthogonal coding provides a neural format capable of maintaining multiple candidate goal outcomes in parallel. This organization contrasts with classical latent-state formulations that infer a single hidden context, and instead aligns with frameworks that permit parallel evaluation of multiple potential outcomes^41,52,53^. Such parallel estimates may facilitate rapid behavioral reorganization when contingencies change, as observed following omission or relocation of reward sites.

Goal-associated representations in the NAc are distinct from those reported in the orbitofrontal cortex (OFC)^24^. Whereas OFC activity reflects the allocentric spatial position of the currently intended destination even during erroneous navigation, the NAc encodes an abstract distance-to-goal signal tied to ongoing movement and incorporates latent representations of previously rewarded locations. This division of labor suggests complementary roles, with OFC representing the selected target and NAc supporting tracking progress toward multiple plausible goal options during navigation.

Although the NAc receives inputs from the hippocampus^54–56^, the persistence of distance-to-goal coding in the NAc during hippocampal and medial entorhinal silencing suggests that it is not solely inherited from allocentric maps. Instead, it likely reflects a transformation within striatal circuits that depends on dopaminergic signaling, integrating information about spatial progress with behavioral relevance and expected outcomes.

How the NAc computes a normalized distance-to-goal remains an open question. Because our experiments were conducted under minimal lighting and did not rely on salient sensory cues, this estimation is likely supported by proprioceptive feedback and path-integration mechanisms rather than by landmark-based calculation. Consistent with this interpretation, the decoded signal more accurately tracked path distance than Euclidean distance in the two-dimensional maze, indicating that it reflects behaviorally accumulated progress along the taken trajectory. Such computations may be implemented within the NAc and local striatal circuits, potentially through interactions with brainstem locomotor pathways^57^. In addition, deriving proportional proximity normalized by total journey length requires an estimate of the target goal location before navigation begins. Given that the OFC provides a major source of inputs to the NAc^58–60^, goal-related signals conveyed by the OFC may play a pivotal role in anchoring distance-to-goal computations to a selected destination.

Together, these results reveal a spatially normalized, goal-invariant state variable in the NAc, establishing a direct neural bridge between cognitive maps and reinforcement learning over physical space. This framework recasts dopamine–striatal function in navigation as operating over orthogonal spatial state variables that simultaneously represent current goals and latent memories of previously rewarded locations. By unifying spatial mapping with value-based learning, this architecture provides a flexible substrate for adaptive long-range goal-directed behavior and suggests a general organizing principle for neural state representations in sequential decision-making.

## Methods

### Subjects

All experiments were approved by the local authorities (RP Darmstadt, protocols F126/1009, F126/1026, and F126/2005) in accordance with the European Convention for the Protection of Vertebrate Animals used for Experimental and Other Scientific Purposes. Fifty male Long-Evans rats weighing 400–550 g (aged 3–6 months) were housed individually in Plexiglass cages (45 × 35 × 40 cm; Tecniplast, GR1800) and maintained on a 12-h light/dark cycle. Behavioral experiments were performed during the dark phase. Six rats were implanted with a Neuropixels1.0^42^ probe, and seven rats were implanted with a Neuropixels 2.0^43^ probe into the NAc. Six rats were injected with AAVs to express hM4Di^45^ in the hippocampus and medial entorhinal cortex. Five rats were injected with AAVs to express ChR2 in DA neurons in the VTA, two of which were further implanted with a tetrode drive and an optic fiber into the VTA, and three were used for histological examination to quantify ChR2 expression in VTA DA neurons. Four rats were injected with AAVs to express stGtACR2^61^ in DA neurons in the bilateral VTA and were implanted with optic fibers. Five rats were injected with the AAV to express dLight1.2^62^ in the NAc and were implanted with an optic fiber. Six rats were injected with AAVs to express eOPN3^51^ in VTA DA neurons, four of which received bilaterally and two unilaterally into the NAc together with a tetrode drive combined with an optic fiber. Four rats were implanted with a Neuropixels 1.0 probe and six rats were implanted with a Neuropixels 2.0 probe into the VTA. Ten rats were implanted with a tetrode microdrive into the hippocampal CA1. Some rats were included in multiple experimental conditions; rat 721 underwent simultaneous recordings in the NAc and VTA; rats 746 and 753 were used for combined NAc–VTA recordings together with hM4Di-mediated hippocampal inhibition; rats 713, 741 and 742 were used for NAc recordings with hippocampal hM4Di inhibition; and rat 754 underwent VTA recordings combined with hippocampal hM4Di inhibition. The animals were water-restricted with unlimited access to food and kept at 90% of their free-feeding body weight throughout the experiment. No statistical method was used to predetermine the sample size.

### Surgery, virus injection, and drive implantation

All surgeries were conducted under anesthesia induced by isoflurane (5% induction concentration, 0.5–2% for maintenance, adjusted according to physiological monitoring). For analgesia, Buprenovet (Buprenorphine, 0.06 mg/mL; WdT) was administered by subcutaneous injection, followed by local intracutaneous application of either Bupivacain (Bupivacain hydrochloride, 0.5 mg/mL; Jenapharm) or Ropivacain (Ropivacain hydrochloride, 2 mg/mL; Fresenius Kabi) into the scalp. The rats were subsequently placed in a Kopf stereotaxic frame, and an incision was made in the scalp to expose the skull. After horizontal alignment, anchor screws were implanted, and craniotomies were made for electrodes (microdrive or Neuropixels), optic fibers, and/or AAV injection. The electrodes and the optic fiber were fixed to the anchor screws with dental cement. Two screws above the cerebellum were connected to the electrode’s ground. All animals received analgesics (Metacam, 2 mg/mL Meloxicam; Boehringer Ingelheim) and antibiotics (Baytril, 25 mg/mL Enrofloxacin; Bayer) for at least 3 days post-surgery.

Simultaneous tetrode recordings and optogenetic manipulations were performed using a custom opto-tetrode microdrive equipped with nine independently adjustable tetrodes (one for reference) as well as an optic fiber (FP200URT, Thorlabs) or lambda-B fiber (OptgeniX, Custom order; NA 0.39, tip length 1.5 mm). The tetrodes were made from 17 µm polyimide-coated platinum-iridium (90–10%; California Fine Wire, plated with gold to impedances below 150 kΩ at 1 kHz). The tetrode bundle consisted of 30-gauge stainless steel cannula, soldered together in a doughnut shape. Microdrive implantation was performed at least 3 weeks after the AAV injection to allow the animals to recover. In two rats, the microdrive was implanted into the right VTA using the following stereotaxic coordinates: anterior–posterior (AP), −5.0 to −6.0 mm; medial–lateral (ML), 0.9 to 1.9 mm, with a 5° angle toward the midline in the coronal plane. The microdrive was initially lowered to a dorsoventral (DV) depth of 4.0 mm from the cortical surface and subsequently advanced to final recording depths of 8.0–8.7 mm. The optic fiber was positioned 0.5–1.0 mm above the tetrode tips. In the remaining two rats, the microdrive targeted the right NAc at the following coordinates: AP, 1.0–2.0 mm; ML, 1.0–2.0 mm. Tetrodes were initially implanted at a DV depth of 4.0 mm from the cortical surface and were gradually lowered to final depths of 7.0–8.0 mm. The AAV injections were conducted at an infusion rate of 100 nl/min using a 10 μl NanoFil syringe and a 33-gauge bevelled metal needle (World Precision Instruments). After the injection was completed, the needle was left in place for 10 min. A volume of 500 nl was injected at each site. The custom microdrive was implanted (see above). The experiments started at least 4 weeks after the virus injection.

For Neuropixels recordings, probes were implanted in the right hemisphere targeting either the VTA (AP, −5.5 mm; ML, 1.4 mm; DV, 8.8 mm) or the NAc (AP, 1.8 mm; ML, 1.5 or 1.8 mm; DV, 8.0 mm). For DREADD-mediated inhibition of the HPC and MEC, AAV8-hSyn-hM4Di was injected bilaterally at multiple sites. For the HPC, injections were made at the following coordinates relative to bregma (in mm): AP −2.5, ML 1.5, DV 2.5; AP −3.5, ML 2.5, DV 2.5; AP −4.5, ML 3.5, DV 2.5; and AP −5.5, ML 4.5, at five DV depths (3.0, 4.0, 5.0, 6.0, and 7.0) using a 5° angle toward the midline. For the MEC, injections were made at 0.8 mm anterior to the edge of the transverse sinus at ML 4.5 and two DV depths (3.5 and 2.5) relative to the cortical surface, using a 25° angle relative to the anterior–posterior axis. For optogenetic identification of DA neurons, AAV2/9-rTH-Cre (Addgene #107788) and AAV2/5-EF1α-DIO-hChR2-mCherry (Addgene #20297) were injected into four sites in the right ventral tegmental area (VTA) at the following coordinates (AP, ML, DV in mm): −5.1, 1.5, 8.5; −5.1, 2.4, 8.5; −5.6, 1.4, 8.0; and −5.6, 2.0, 8.0, using a 5° angle toward the midline. For fiber photometry measurements of DA concentration, AAV2/9-hSyn-dLight1.2 (Addgene #111068)^62^ was injected into the right NAc at the coordinates of AP, 1.8; ML, 1.8; DV 7.3 in mm. An optical fiber (FP400URT, Thorlabs) was implanted with its tip positioned approximately 500 μm above the injection site (AP, 1.8; ML 1.8; DV, 6.8 in mm). For optogenetic inactivation of DA neurons in the VTA, AAV2/9-rTH-Cre and AAV2/1-hSyn-SIO-stGtACR2-FusionRed (Addgene #105677)^61^ were injected into two sites in each hemisphere of the VTA at the following stereotaxic coordinates (AP, ML, DV in mm): −5.1, 1.7, 8.5 and −5.6, 1.4, 8.0, using a 5° angle toward the midline. To inactivate a larger population of neurons, lambda-B optical fibers (OptgeniX; custom order) with a 1.5-mm tapered tip were used. These fibers were implanted bilaterally with their tips positioned in the VTA at AP −5.5 mm, ML 1.4 mm, and DV 8.5 mm, using a 5° angle toward the midline. For optogenetic inhibition of DA release in the NAc, AAV2/9-rTH-Cre and AAV2/1-CaMKII-eOPN3-mScarlet (Addgene #125712)^51^ were co-injected into two sites in the VTA at the following stereotaxic coordinates (AP, ML, DV in mm): −5.1, 1.7, 8.5 and −5.6, 1.4, 8.0, using a 5° angle toward the midline. For experiments involving simultaneous recordings of neural activity in the NAc under optogenetic manipulation, viral injections were performed unilaterally, and a custom tetrode microdrive equipped with an optical fiber was implanted in the same region. For the behavioral experiments shown in Fig. 3f, viral injection and implantation of lambda-B optical fibers were performed bilaterally in the NAc with their tips positioned at AP 1.8 mm, ML ±1.5 mm, and DV 7.8 mm. For recording from the hippocampus, a 28-tetrode microdrive was implanted bilaterally at AP −4.0 mm, ML 3.2 mm with the depth adjusted until observing clearly separated clusters with sharp-wave ripples.

### Drug administration and optogenetic manipulation

For hM4Di-mediated inhibition, deschloroclozapine (DCZ)^46^ was used as the DREADD ligand. DCZ was administered subcutaneously at a fixed dose of 50 µg per animal, 1 h before the experiment. Based on body weight (400–550 g), this corresponds to an effective dose of approximately 0.09–0.13 mg/kg. To assess drug effects, we also conducted saline-injection sessions as well as no-injection control sessions within the same animals.

For eOPN3 activation, green laser pulses (500 ms duration at 0.1 Hz) were delivered from a 532-nm DPSS laser (Shanghai Laser & Optics Century, GL532T3-100) for approximately 5 min. Laser power at the fiber tip in each hemisphere was ∼10 mW. For GtACR2 activation, continuous blue light was delivered from a 473-nm DPSS laser for approximately 5 min, with laser power at the fiber tip of each hemisphere set to ∼20 mW. For ChR2 activation, blue laser pulses (1, 3, 5, or 10 ms duration at 0.1 Hz) were delivered from a 473-nm DPSS laser (Shanghai Laser & Optics Century, BL473T8-200) while animals were resting on a pedestal. Laser power at the fiber tip was approximately 30 mW.

The onset of laser stimulation was manually triggered at the time of goal change, and the timestamps of all laser pulses were recorded.

### Behavioral tasks

#### Linear maze task

The linear maze was 2 m in length and equipped with 10 identical water-delivery wells on the floor. Each well contained an infrared optical sensor to monitor licking behavior. Behavioral training was conducted in multiple phases. During the initial habituation phase, water rewards were delivered at several wells, and additional rewards were provided at wells that the animal licked to facilitate the association between well licking and reward delivery. In the subsequent training phase, water was delivered only when the animal alternated licking between a fixed pair of goal wells. A lick-duration threshold was then introduced and gradually increased, requiring animals to maintain licking at the correct well for a fixed duration (1.0 – 1.5 s) before reward delivery, thereby reinforcing goal confidence. In the final phase, the identity of the goal-well pairs was changed multiple times within each daily session.

Recording and behavioral experiments began once animals were able to adapt their behavior to goal changes more than three times per day. During the initial trials following the introduction of a new goal pair, water was prefilled at the goal wells and LEDs beneath all wells were illuminated to signal the goal change. In subsequent trials, animals were required to visit and lick the correct well for a fixed duration to receive water delivery. The animal’s position and head direction were tracked using two colored LEDs mounted on the animal’s head at a sampling rate of 20 Hz.

The start of each journey was defined as the position at which the animal’s running speed first exceeded 20 cm/s, and the goal location was defined as the position at which the animal first licked the correct goal well. Trials in which the animal licked an incorrect well for more than 0.5 s were classified as error trials. All recordings were performed under minimal lighting conditions, with no direct light sources in the recording space and only weak stray light from computer monitors behind curtains, to prevent cue-guided navigation.

#### Two-dimensional maze task

The open-field square maze measured 1.5 × 1.5 m and was equipped with 25 water-delivery wells embedded in the floor, arranged in a 5 × 5 grid with equal spacing. Each well contained an infrared optical sensor to monitor licking behavior. Behavioral training was conducted in multiple stages. First, animals were placed in a small walled compartment containing four wells, where they were allowed to explore the environment. Licking at any well was paired with water delivery, facilitating the association between well licking and reward. Next, an alternation rule was introduced within the same four-well compartment. One well was designated as a fixed home well. Animals were first required to lick the home well, after which an LED beneath one of the remaining three wells was illuminated. Animals were then required to visit and lick the illuminated well to receive a water reward. A lick-duration threshold was introduced, requiring animals to maintain licking at the correct well for a fixed duration (1.0 s) before reward delivery. Following successful completion, animals were again required to return to the home well. This sequence was repeated with a fixed home well and randomly designated cued wells. After animals achieved a performance criterion of >80% correct responses, the compartment size was expanded to include 10 wells and subsequently to the full 25-well arena. The home well was changed between daily sessions but remained fixed within each session. Once animals learned the task, recording sessions commenced. Animal position and head direction were tracked using two colored LEDs mounted on the animal’s head. The beginning of each journey was defined based on a running-speed threshold together with displacement from the start well.

#### Object location task

To assess the behavioral impact of DREADD-mediated neural suppression, rats were tested in an object location task. The task was conducted in an open-field arena measuring 50 × 50 cm. Animals were habituated to the empty arena for two consecutive days before the start of object location testing. Multiple objects were fabricated using a 3D printer from the same black resin (Black Resin V4, Formlabs) and were matched in size but differed in shape (cube or triangular prism). Object identity was randomly assigned for each experimental session. For the experimental procedure, saline or the DREADD agonist deschloroclozapine (DCZ; 50 µg) was administered subcutaneously 1 h before behavioral sessions. Animals were then placed in the arena containing two identical objects and allowed to freely explore for 15 min (familiarization phase). After a 1-h retention interval in the home cage, animals were returned to the same arena for a 15-min test phase, during which one of the objects was relocated to a novel position. To assess exploration bias between the two objects, we analyzed only periods of object exploration during the test phase, defined as times when the animal’s position was within an 8-cm radius of either object. The analysis was segmented minute-by-minute based on cumulative object exploration time. Object location memory performance was quantified as the proportion of total exploration time directed toward the relocated object and was compared across DCZ-treated and saline-injected control sessions.

#### Linear maze task with reward omission

Reward omission tests were conducted after rats had been fully trained on the linear maze task. At the start of each behavioral session, animals navigated between a given pair of goal wells. After performance stabilized, defined as at least five consecutive correct trials, one well of the goal pair near the center of the maze was changed to a new location, and reward delivery was resumed until the animal again achieved five consecutive correct trials. Once this performance criterion was met, reward delivery at the newly assigned goal well (but not at the track-end well) was suspended for approximately 2 min. During the omission period, animals typically ceased licking at the reward-omitted well and began exploring other wells. For behavioral analysis, licking events lasting longer than 0.5 s at wells outside the current goal pair were counted. Licked wells were ranked according to their distance from the well that had been rewarded in the previous block but was changed to a new location in the current block. The resulting distributions were compared against a chance level of 13%, assuming uniform selection among all non-goal wells, using a binomial test.

To test the impact of VTA dopaminergic inputs to the NAc on the animal’s ability to execute memory-guided search for previously rewarded locations, reward omission experiments were performed in rats expressing the inhibitory opsin eOPN3 selectively in VTA dopamine neurons via Cre recombinase driven by the tyrosine hydroxylase promoter. Green laser illumination (532 nm; 500 ms duration at 0.1 Hz) was delivered to the NAc to suppress dopaminergic terminal activity. To ensure comparable baseline behavior across animals, omission experiments were initiated only after rats reliably revisited the previous goal location during an initial omission session without any manipulation (otherwise excluded). Subsequent omission tests were conducted across four consecutive days with or without optogenetic manipulation.

### Histological processing

Upon completion of experiments, animals were deeply anesthetized with sodium pentobarbital and transcardially perfused with saline followed by 10% formalin. Brains were extracted and post-fixed in formalin at 4 °C for at least 72 h. Fixed brains were then frozen and sectioned coronally at 30 μm thickness using a cryostat. To verify the locations of tetrodes, optical fibers, and Neuropixels probes, coronal or sagittal sections encompassing the implantation sites were directly mounted onto glass slides, stained with cresyl violet, and coverslipped using mounting medium (HISTOMOUNT, Invitrogen). To confirm AAV-mediated protein expression, sections were immunostained to enhance fluorescence signals of hM4Di-mCherry, ChR2-mCherry, GtACR2-FusionRed, eOPN3-mScarlet, and dLight1.2. Sections were first incubated for 30 min in blocking buffer consisting of 4% normal goat serum and 2% bovine serum albumin in 10× TBS (pH 7.6).

For co-staining of ChR2-mCherry and tyrosine hydroxylase, sections were incubated for 36 h at 4 °C with a mouse anti–tyrosine hydroxylase antibody (1:250; Sigma, T2928) and a rabbit anti–red fluorescent protein antibody (1:500; Abcam, ab62341). Sections were then washed for 30 min in TBS containing 0.2% Triton X-100 and incubated for 4 h at 4 °C with Alexa Fluor 488–conjugated goat anti-mouse antibody (Invitrogen, A11001) and Alexa Fluor 594–conjugated donkey anti-rabbit antibody (1:250; Invitrogen, A21207).

After additional washes in TBS containing 0.2% Triton X-100, sections were mounted on glass slides and coverslipped using ProLong Gold antifade reagent with DAPI (Invitrogen, P36931). Immunostaining of hM4Di-mCherry, GtACR2-FusionRed, and eOPN3-mScarlet was performed using the same primary and secondary antibodies as for ChR2-mCherry. For dLight1.2 staining, sections were incubated with a chicken anti-GFP antibody (1:200; Invitrogen, A10262) followed by a goat anti-chicken Alexa Fluor 594 secondary antibody (1:500; Invitrogen, A32759). All images were acquired using a slide scanner (Zeiss Axio Scan.Z1).

ChR2-expressing neurons were identified based on mCherry fluorescence, and co-localization with tyrosine hydroxylase immunoreactivity was quantified. Across four rats, 87% of ChR2-expressing neurons (786 of 899 cells) were positive for tyrosine hydroxylase, confirming that a majority of the ChR2-expressing neurons were dopaminergic, as reported previously^63^.

### Spike sorting and cell classification

Data processing and analyses were performed using MATLAB (MathWorks) and Python. For tetrode microdrive recordings, neural signals were acquired and amplified using RHD2164 headstages (Intan Technologies) at a sampling rate of 15 kHz with the Open Ephys acquisition system. For recordings with Neuropixels probes, neural signals were acquired using a PXIe acquisition module (IMEC) housed in an NI PXIe-1071 chassis (National Instruments), also using the Open Ephys acquisition system. Signals were band-pass filtered between 0.6 and 6 kHz, and spikes were detected and clustered using Kilosort (https://github.com/cortex-lab/KiloSort)^64,65^. For tetrode recordings, Kilosort1 was used with individual tetrodes grouped via the kcoords parameter, and the noise parameter controlling the fraction of noise templates shared across channel groups was set to 0.01. Resulting clusters were manually curated based on autocorrelograms and waveform features in principal component space, with multiunit activity and noise discarded to obtain well-isolated single units. Neurons with mean firing rates below 0.5 Hz were excluded from further analyses. Firing rates were estimated by convolving spike times using a Gaussian kernel with a bandwidth of 100 ms.

To identify ChR2-expressing neurons in opto-tetrode microdrive recordings, spiking activity was recorded during delivery of blue laser pulses (1, 3, 5, or 10 ms duration at 0.1 Hz) to the recording region. For each neuron, firing rates were converted to z-scores using the mean and standard deviation computed from a 50-ms baseline window preceding laser onset. Neurons exhibiting firing rates exceeding a z-score of 10 within a 10-ms window following laser onset were classified as ChR2-expressing.

### Uniform Manifold Approximation and Projection

To examine whether NAc population activity during repeated goal-directed journeys is organized in a conjunctive or disentangled manner, we visualized the geometric structure of neural population activity using Uniform Manifold Approximation and Projection (UMAP)^44^ embedded in three dimensions. UMAP was performed using the umap-learn Python library^44^ with standard parameters (n_neighbors = 30, min_dist = 0.1, n_components = 3) based on Euclidean distance.

Neural activity was smoothed using a 500 ms moving window. Each journey epoch was segmented into running and stationary phases: from motion onset to the first lick at the goal well (running), and from the first lick at the goal well to the subsequent motion onset (stationary). Each phase was temporally normalized and divided into 20 equal time bins, yielding a total of 40 bins per journey epoch. For dimensionality-reduction analyses, firing rates of each neuron were standardized by z-scoring across all epochs. The resulting three-dimensional UMAP embeddings were visualized for both phases using graded colors to represent the temporal evolution of neural population activity.

For quantitative analyses, persistent homology (PH) was computed on the three-dimensional UMAP embeddings described above using the ripser Python library for homology groups H0 and H1^66,67^. H0 persistence was used to verify that population trajectories formed a single continuous structure. Quantitative comparisons focused on the persistence (death-birth) of finite H1 features. For each behavioral block and journey condition, trials were randomly subsampled to equalize sample numbers across conditions. Statistical significance was assessed relative to shuffle controls in which coordinates were independently permuted within each UMAP dimension. This procedure was repeated 500 times to generate null distributions for comparison with real data.

To summarize ring-shaped topology across blocks, the largest finite H1 persistence (H1_max) was compared against shuffle-derived null distributions. For each inbound and outbound trial group, an H1_max percentile was defined as the proportion of shuffle realizations (500 coordinate shuffles) for which the real H1_max exceeded the shuffled value. To obtain block-level statistics, inbound and outbound trial groups were examined, and the smaller of the two was taken within each block. This approach ensured that ring structure was consistently present across both movement directions rather than driven by a single trajectory subset. The proportion of blocks exceeding the 97.5% shuffle percentile was evaluated using one-sided exact binomial tests relative to the null probability expected from shuffle controls. To quantify ring multiplicity, the number of significant H1 persistence bars per trial group was defined as the count of finite H1 lifetimes exceeding the 97.5% shuffle percentile. For block-level summaries, ring counts from inbound and outbound trial groups were averaged to avoid inflation of ring numbers caused by directional duplication. The resulting distributions of mean ring counts across blocks were visualized as histograms. The prevalence of single-ring blocks was assessed using a one-sided exact binomial test, evaluating whether the observed proportion of blocks with exactly one significant ring exceeded the null probability expected under an equal-likelihood distribution across ring-count categories.

### Support Vector Regression

Support vector regression (SVR) was used to decode the goal proximity from NAc population activity. Four candidate proximity variables were evaluated: absolute distance to the goal, elapsed time, proportional distance (distance normalized by total path length), and proportional time (time normalized by total trial duration). Neural activity was aligned to the animal’s trajectory from movement onset to goal arrival and divided into 20 equally spaced position bins. Only correct trials were included. For each neuron, firing rates were normalized using min–max scaling to the range [0, 1] based on activity across the entire recording session. To assess generalization across spatial contexts, decoding was performed across four behavioral blocks with distinct goal locations. For each daily session, SVR models were trained on data from two blocks and tested on the remaining two blocks, using all six possible train–test combinations (i.e., all combinations of 2 blocks chosen from 4). Both outbound and inbound trajectories were included in the training and testing datasets.

Decoding was performed using a linear SVR implemented in Python (scikit-learn) and MATLAB (fitrsvm, Statistics and Machine Learning Toolbox, MathWorks). Neural population firing rates served as input features, and the behavioral proximity variable served as the target output. Predictions were generated on held-out test trials on a trial-by-trial basis, yielding a predicted proximity value for each spatial bin. Decoding performance was then quantified using the scaled mean squared error (SMSE), defined as the mean squared prediction error normalized by the power of the ground-truth signal:

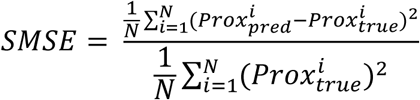

where 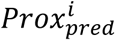 denotes the SVR-predicted proximity value at the spatial bin 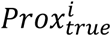 denotes the corresponding ground-truth proximity value, and 𝑁 is the total number of spatial bins (20). This normalization expresses prediction error relative to the intrinsic power of the behavioral variables, allowing direct comparison of decoding performance across different proximity measures with distinct units and dynamic ranges. Lower SMSE values indicate more accurate decoding, with values approaching zero reflecting near-perfect prediction. SMSE values were computed for each trial and used for statistical analyses.

Fast and slow running trials were defined to dissociate time- and distance-based coding. Trials were classified solely based on running-speed profiles. Trials were labeled as fast when running speed remained above 30 cm/s after exceeding this threshold and until reaching the goal, and as slow when running speed dropped below 10 cm/s in the middle of the journeys. Decoding performance for fast and slow trials was evaluated using the same support vector regression model within each recording session. For each session, a decoder for proportional goal distance was trained on the full population of simultaneously recorded NAc neurons and then applied separately to fast and slow trial subsets. Statistical comparisons of decoding error were performed using a two-way repeated-measures ANOVA with factors of speed (fast vs. slow) and axis (distance vs. time) to assess whether decoding performance reflected distance rather than elapsed time.

#### Goal distance decoding in the two-dimensional navigation task

For the two-dimensional navigation task, proportional distance to the goal was computed as the cumulative path distance from the animal’s instantaneous position to the target well along its trajectory, divided by the total path distance from journey onset to that target. Because starting positions varied across trials (unlike in the linear maze), normalization was performed separately for each trial. Neural activity during navigation was segmented into 20 equally spaced bins between journey onset and goal arrival. Decoding analyses were performed using a leave-one-trial-out cross-validation scheme. SVR decoders were trained on home-well–directed trials and subsequently applied to both home-well–directed and random-cue–targeting trials to assess generalization across task conditions.

#### Construction of decoders for current- or previous goals

To assess whether NAc population activity independently represents proximity to the currently targeted goal and to a previously visited goal, we implemented a dual-decoder SVR strategy. Two linear SVR decoders were trained on neural population activity recorded across behavioral blocks. Both decoders were trained using data from the first two blocks, during which animals navigated to distal and proximal goal wells, respectively. In the third block, when animals again targeted the distal well, one decoder was trained to predict proximity to the goal visited in the immediately preceding block (proximal well), whereas the second decoder was trained to predict proximity to the currently targeted goal (distal well). All ramp-up neurons recorded in the NAc were included in the decoding analysis. Trials were pooled across daily sessions and animals. Decoder performance was evaluated using a leave-one-trial-out cross-validation procedure by comparing predicted and true proximity values on held-out trials. For visualization and population geometry analyses in Fig. 4f, linear SVR decoders were additionally trained on pooled trials from the same blocks without cross-validation.

#### Angle calculation between decoder subspaces

To characterize the population-level relationship between information subspaces for current and previous goals, we quantified the angle between their regression vectors in neural population space. The two decoders for proportional goal distance optimized for either the current or the previous goal were constructed from the same NAc neural population activity.

To estimate the statistical distribution of decoder angles, a bootstrap resampling procedure was performed. Trials were resampled with replacement, and the two decoders for the current and the previous goals were retrained on each resampled dataset using the same procedure described above. The angle between the resulting regression vectors was computed for each resample, yielding a bootstrap distribution of angles between two decoding axes. A total of 1,000 bootstrap resamples were performed. Bootstrap resampling was used solely to visualize the variance of decoder angles and was not used for statistical inference or hypothesis testing.

### Cell classification

#### Hierarchical clustering analysis

To characterize population-level activity patterns during navigation, firing rates of individual neurons were z-scored and averaged across trials along a distance-normalized journey, defined from motion initiation to the first lick at the goal well. Each journey was divided into 20 equally spaced position bins. For each neuron, an activity vector was constructed from the resulting z-scored firing rates across bins. Agglomerative hierarchical clustering was performed on these activity vectors using correlation distance and average linkage. The number of clusters was determined using the elbow method applied to the within-cluster sum of squares from K-means clustering. Clustering was used descriptively to group neurons with similar temporal activity profiles.

#### Selection of ramp-up and ramp-down firing neurons

For analyses involving unilateral dopaminergic inhibition in Fig. 3i, NAc neurons were classified based on task-related modulation of firing near the goal location. Spike rates were aligned to normalized position from journey onset to goal arrival. Each trial was divided into 20 equally spaced position bins. For each neuron, firing rates were z-scored. Activity in an early segment of the trajectory (bins 1–10) was compared with activity in a late segment corresponding to the goal region (bins 19–20). For each trial, the mean firing rate in the late segment was subtracted from that in the early segment, yielding a trial-wise difference value. To assess whether firing activity near the goal differed consistently from early activity, a paired nonparametric Wilcoxon signed-rank test was applied to the trial-wise differences for each neuron (p < 0.05). Neurons exhibiting a significant increase in activity in the late segment were classified as ramp-up neurons, whereas neurons exhibiting a significant decrease were classified as ramp-down neurons.

### Cosine similarity analysis

To quantify the consistency of population activity patterns across blocks, we computed cosine similarity between block-level population activity vectors for each cluster. For each block, neural activity was first averaged across neurons within each cluster to obtain a cluster-level firing rate profile. Activity profiles were computed separately for inbound, rest, outbound, and subsequent rest epochs, and concatenated to form a one-dimensional population activity vector for each block. Prior to similarity computation, each block-level vector was z-scored to remove differences in overall firing rate magnitude across blocks. Cosine similarity was then calculated between all pairs of block-level population activity vectors within each cluster. This metric captures the similarity of activity patterns while being insensitive to absolute firing rate scaling.

### DA measurement by *in vivo* fiber photometry

To measure DA concentration in the NAc of behaving animals, fiber photometry recordings were performed using a wireless photometry system (TeleFipho, Amuza Inc.). An optical fiber (FP400URT, Thorlabs; length, 25 mm) was implanted above the NAc. A blue LED (bandpass-filtered; 445–490 nm) was used to excite the genetically encoded dopamine sensor dLight1.2^62^, and emitted fluorescence was detected by an internal photodiode optimized for green light (bandpass-filtered; 500–550 nm). Fluorescence signals were amplified by an internal DC amplifier and transmitted wirelessly as 16-bit digital signals to a receiver, allowing real-time monitoring using TeleFipho software. Prior to each daily recording session, LED power was adjusted to achieve an optimal signal amplitude while avoiding saturation. The analog output from the wireless receiver was acquired by the OpenEphys acquisition system and synchronized with behavioral video-tracking data.

Signal baseline was defined as the mean within a 1 s window following the final lick at the goal well in the preceding trial. This baseline was subtracted from the raw fluorescence signal of the subsequent trial. Baseline-corrected signals were then normalized using min–max scaling, with scaling parameters computed from trials within the same goal block.

### Statistical analysis

Statistical analyses were conducted using the Python SciPy library and MATLAB (2021a) statistical toolbox (MathWorks). All statistical tests were two-sided and non-parametric unless stated otherwise.

