## Extended Data Figures for "A multiplexed striatal architecture for generalized spatial goal progress"

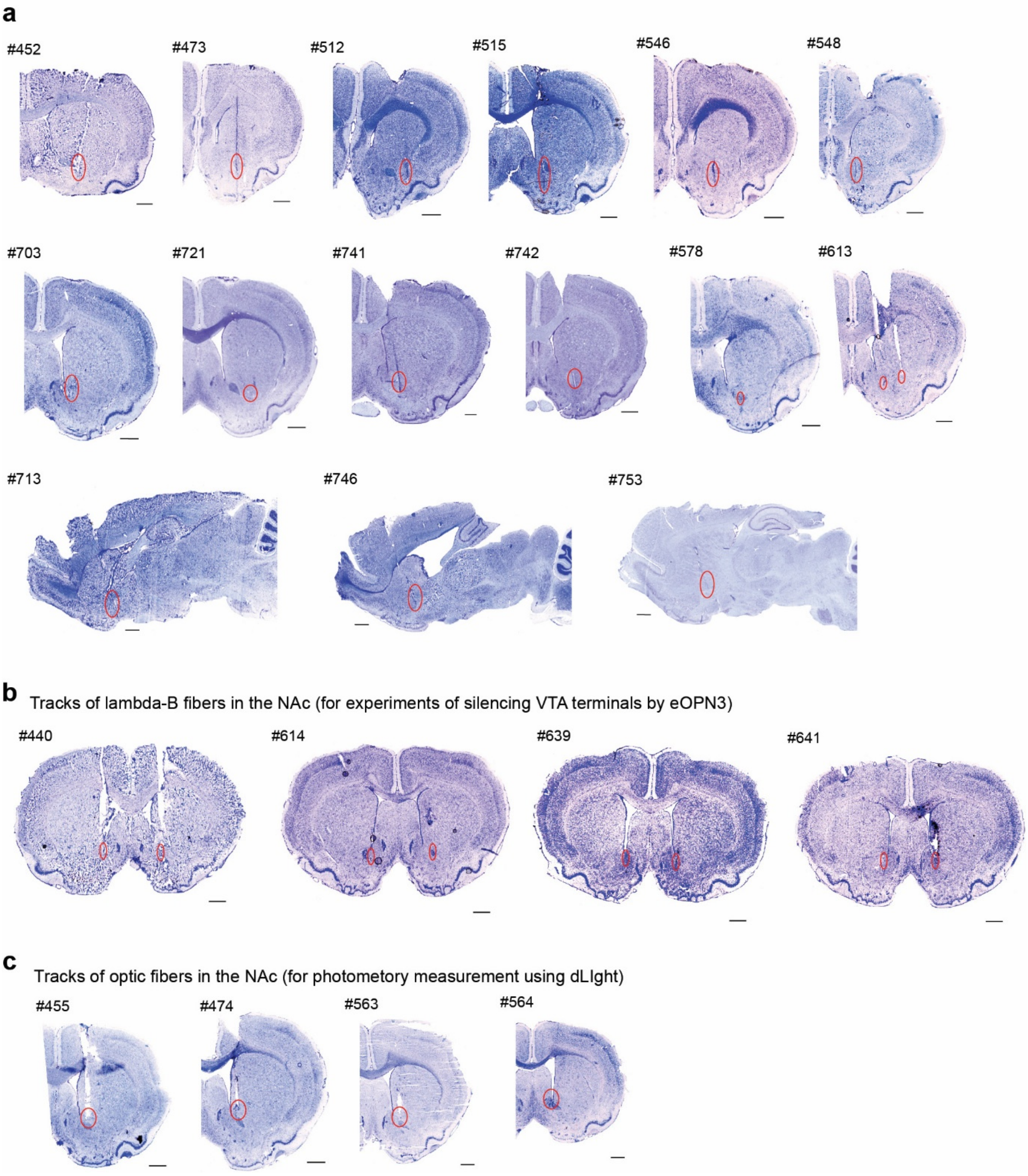


**Extended Data Figure 1. Histological verification of electrode or optic fiber implantation sites in the NAc.**

**a,** Nissl-stained coronal sections showing recording tracks of Neuropixels probes (animals #452, 473, 512, 515, 546, 548, 703, 721, 741, and 742) and tetrodes (animals #578 and 613) implanted in the NAc. Additional sagittal sections illustrate Neuropixels probe tracks targeting the NAc (animals #713, 746, and 753). Red circles indicate the locations of electrode tracks. Scale bar, 1 mm. **b,** Nissl-stained coronal sections showing optical fiber tracks implanted bilaterally in the NAc for eOPN3-mediated silencing of VTA dopaminergic inputs. **c,** Nissl-stained coronal sections showing optical fiber tracks for fiber photometry measurement experiments by using the dopamine sensor dLight1.2

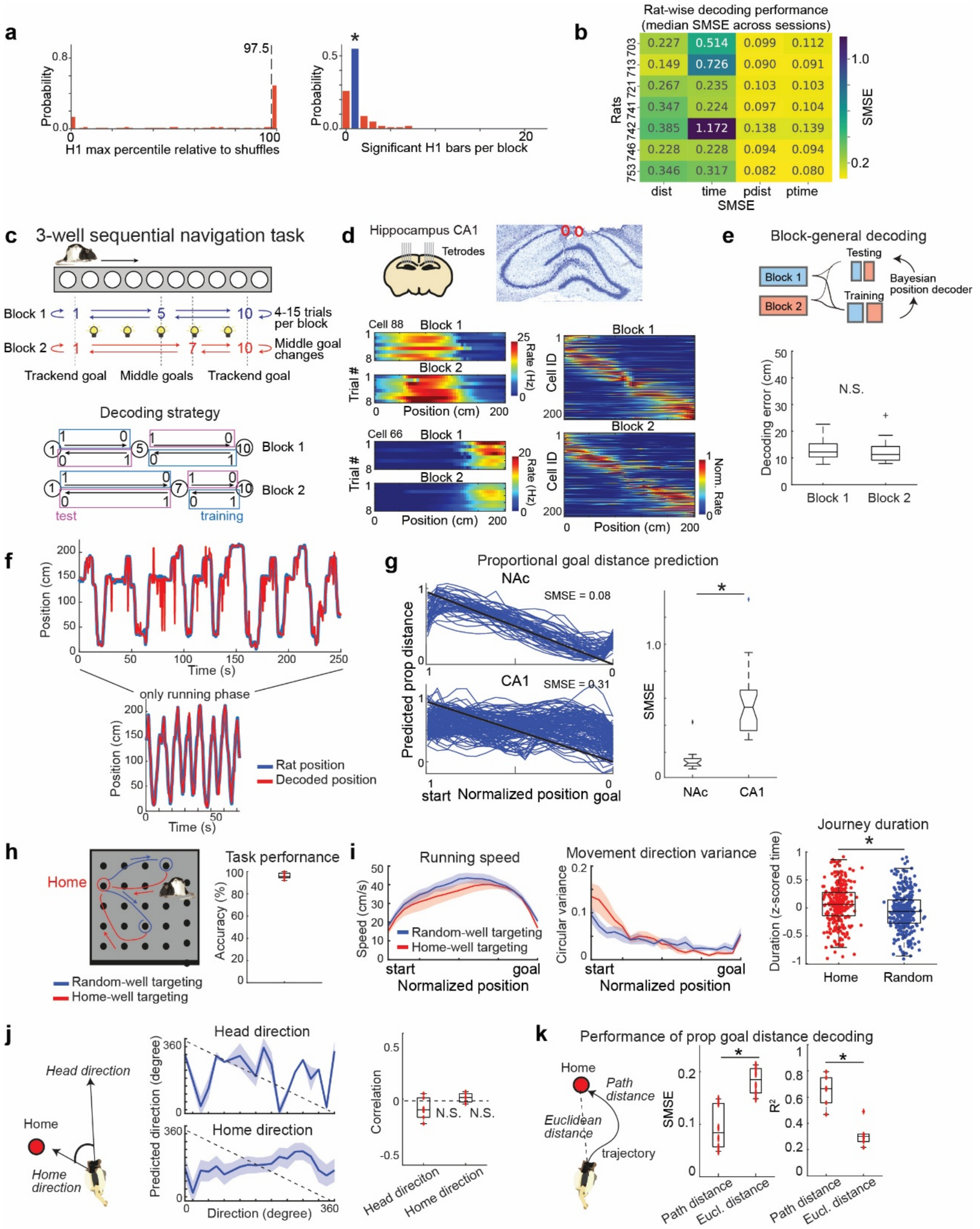


**Extended Data Figure 2. Population activity in the NAc, but not hippocampal CA1, tracks goal proximity that generalizes across start–destination combinations in both one- and two-dimensional tasks.**

**a,** Block-wise consistency of ring-shaped topology on UMAP embeddings. Left, Distribution of H1_max percentiles relative to shuffled datasets (500 coordinate shuffles). The vertical dashed line indicates the 97.5th percentile. For block-level statistics, inbound and outbound trial groups were combined and the smaller percentile value between the two directions was considered. The fraction of blocks exceeding the 97.5th percentile (53/104 blocks, 50.96%)

was significantly higher than expected by chance (binomial test, $p < 0.001$). Right, Distribution of the mean number of significant H1 bars per block. For each inbound and outbound trial group, H1 persistence bars exceeding the 97.5th percentile were counted, and the average of the two directions was computed. Blue bars highlight blocks with a single significant ring. Single-ring blocks were more frequent than zero-ring blocks (57 vs. 27 blocks; binomial test, $p < 0.001$) and more frequent than blocks with multiple rings (57 vs. 20 blocks; $p < 0.001$), indicating a dominant single-ring topology. **b,** Animal-wise decoding performance across behavioral variables. Same analysis as in Fig. 2b, with median SMSE values shown separately for each rat. For each rat, decoding performance was evaluated using SVR and quantified by SMSE. Median SMSE values are shown as a color-coded matrix. SMSEs for proportional distance and proportional time remained consistently low across animals, indicating that decoding performance was robust and not driven by specific animals. **c,** Schematic illustrating the 3-well sequential task. The linear maze was identical to that used in Fig. 1a. In each block, three wells constituted the goal set: two located at the ends of the track and one positioned at the center. After 4–15 trials (including both outbound and inbound runs), the central goal well was changed, requiring the animal to update the goal location. Bottom: Schematic illustrating the decoding strategy. Training and testing data were selected from distinct journey epochs within the session. **d,** Top: Schematic and Nissl-stained coronal section showing tetrode implantation in hippocampal CA1. Bottom left: Color-coded firing-rate maps of two representative CA1 neurons across trials and blocks. Bottom right: Color-coded firing-rate maps of 200 representative CA1 neurons, demonstrating stable spatial coding across blocks. **e,** Assessment of position decoding stability between blocks. The animal's instantaneous position was estimated from CA1 population activity using a Bayesian decoding approach. Data from Blocks 1 and 2 were combined and used for training and testing, with analyses restricted to the running phases of the sessions. Decoding errors did not differ significantly between Blocks 1 and 2, indicating stable spatial representations across blocks. **f,** Plots depicting example trials showing both the animal's position (blue) and the decoded position from CA1 population activity (red). Top: Continuous time across trials. Bottom: Extracted view of the running phases, illustrating particularly accurate decoding during locomotion. **g,** Left: Example trials showing predictions of proportional distance to the goal from NAc and CA1 population activity in the three-well task shown in **c**, aligned to normalized position from start to goal. Predictions from individual trials are shown in blue, with identity lines in black. Right: Box plots of SMSE for proportional goal-distance decoding from NAc and CA1 population activity, demonstrating significantly better decoding performance in NAc compared with CA1 ($p < 0.001$; Wilcoxon rank-sum test; 10 rats, 25 daily sessions). **h,** Schematic of the goal-directed navigation task in the two-dimensional maze, as in Fig. 2g. Right: Task performance quantified as the percentage of correct targeting to the home well (6 sessions from 2 animals). **i,** Comparison of running speed (left), movement direction variance (middle), and journey duration (right) between random-well-targeting and home-well-targeting trials. Overall behavioral profiles were similar across conditions; however, animals exhibited slightly faster and smoother trajectories during random-well-targeting trials, likely due to the cue-guided nature of these runs. **j,** Decoding of head direction and home-well direction from NAc population activity. Support vector regression was used to estimate the relationship between population activity and each directional variable. As shown in the right panel, decoding performance did not differ significantly from zero correlation for either variable ($p > 0.05$; Wilcoxon signed-rank test). **k**, Comparison of proportional goal distance decoding performance based on path-distance versus Euclidean-distance calculations. Left, schematic illustration of differences between path distance and Euclidean distance representations. Middle: Box plots showing SMSE of decoding performance, where decoding based on path distance exhibited significantly lower SMSE than that based on Euclidean distance ($p < 0.05$, Wilcoxon signed-rank test). Right, box plots showing the coefficient of determination ($R^2$). Path-distance–based decoding also produced significantly higher $R^2$ values ($p < 0.05$). The superior fit for path distance indicates that NAc activity reflects cumulative movement along the selected path, consistent with internal computation of self-motion–derived signals.

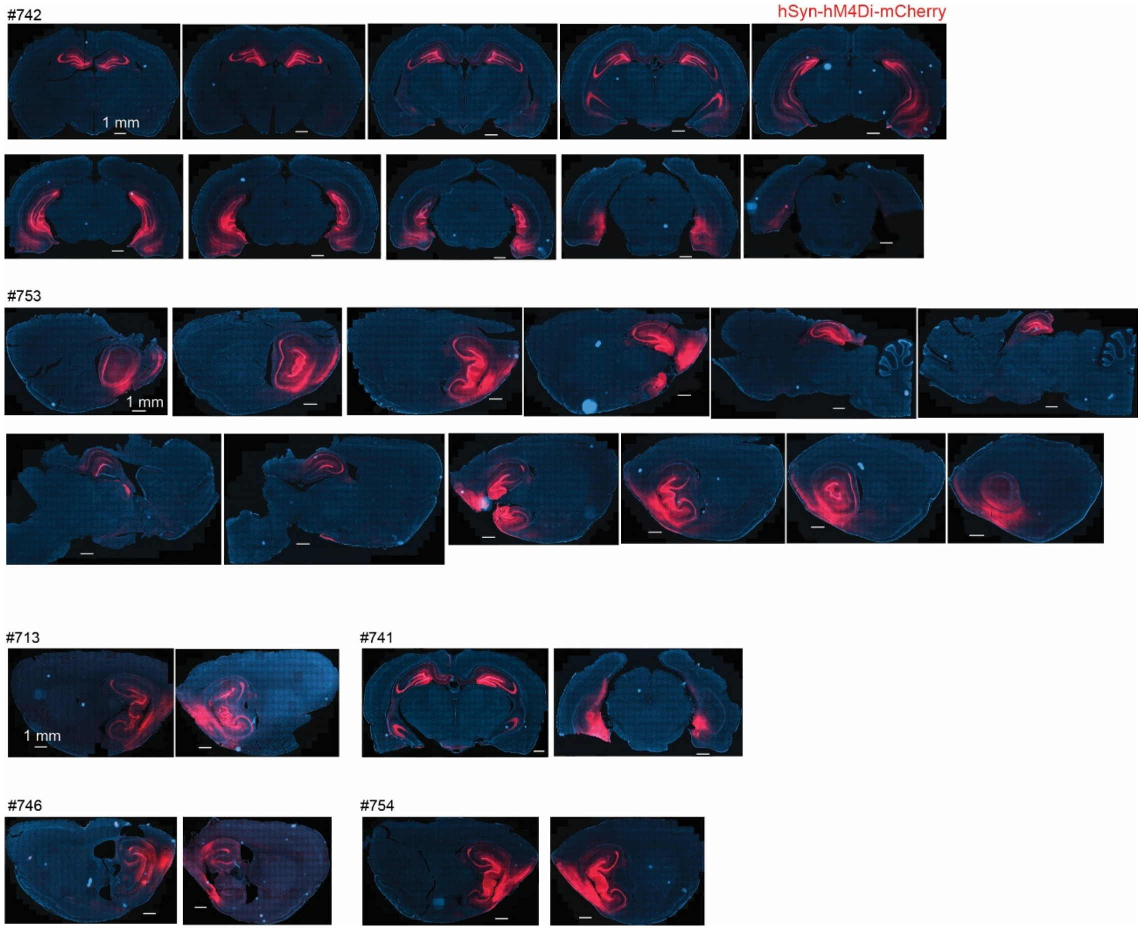


**Extended Data Figure 3. Immunostaining confirming hM4Di expression in the hippocampus and medial entorhinal cortex.**

Coronal sections are shown for animals #741 and #742, and sagittal sections for animals #713, #746, #753 and #754. Red fluorescence indicates hM4Di–mCherry expression detected using an anti-RFP antibody. Scale bar, 1 mm.

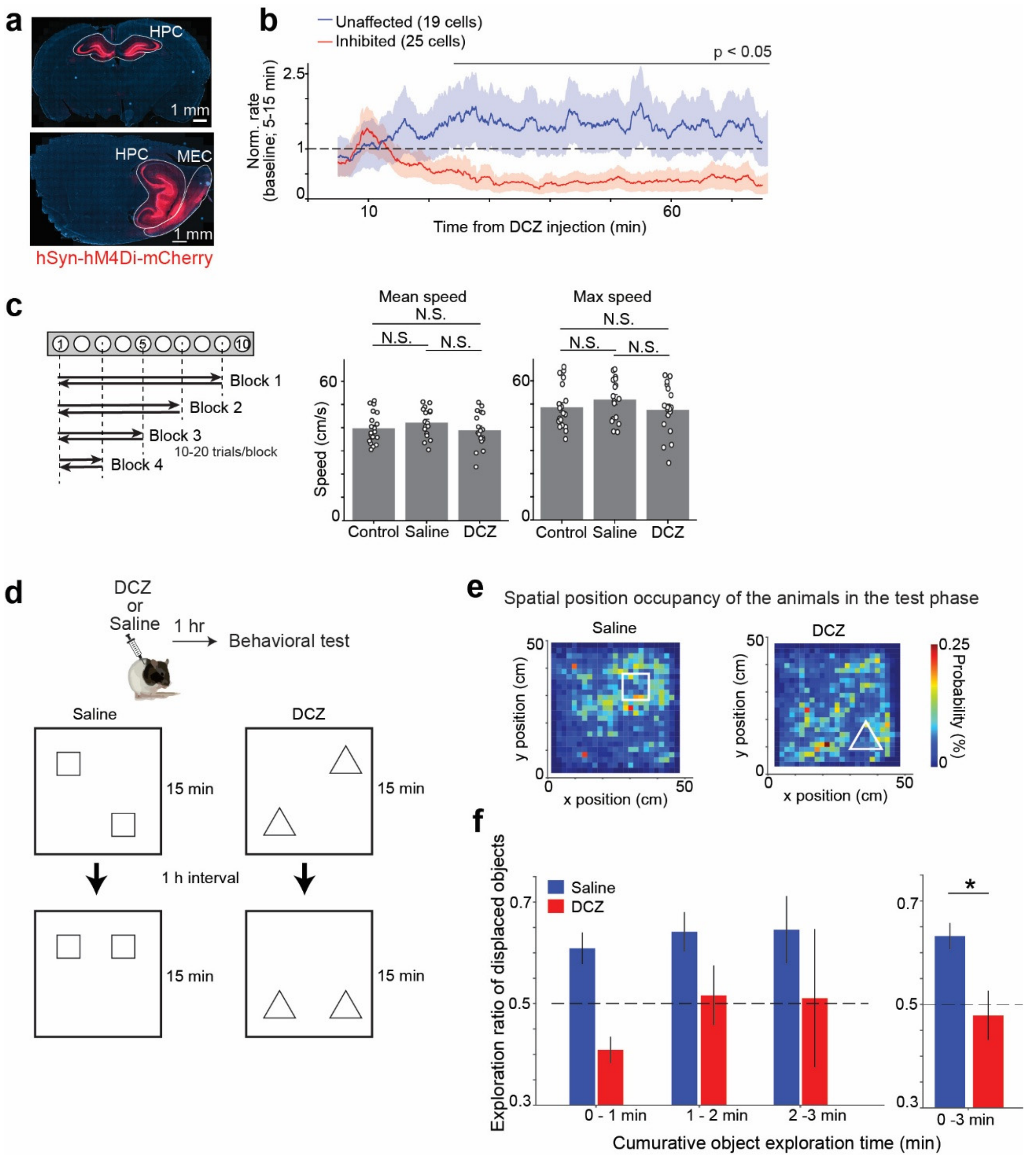


**Extended Data Figure 4. Assessment of DREADD-mediated hippocampal silencing in a hippocampus-dependent object location task.**

**a,** Representative immunostained section showing hM4Di–mCherry expression in the hippocampus (HPC) and medial entorhinal cortex (MEC). Red fluorescence indicates hM4Di expression. Scale bar, 1 mm. **b,** Example recordings from CA1 neurons during DCZ administration. This experiment was performed to validate the DREADD system, and the animal was not included in other experiments. A tetrode microdrive was implanted in hippocampal CA1 of an animal injected with AAV-hSyn-hM4Di-mCherry. After stable CA1 units were obtained, DCZ was administered subcutaneously and firing rates were monitored. The plot shows normalized firing rates, with more than half of the recorded neurons exhibiting a significant reduction in firing within 30 min after DCZ injection. **c,** Assessment of the impact of HPC and MEC silencing on running speed during the navigation task. The animal's running speed was compared across non-injected control, saline, and DCZ-injected sessions. No significant differences were observed in either mean running speed or maximum speed across conditions. **d,** Schematic of the object location memory task. Rats explored two objects in an open-field square arena (50 × 50 cm) during a 15 min habituation phase, followed by a 1 h interval in the home cage. Animals were then returned to the same arena for the test phase, during which one object was relocated. Object identities and locations were counterbalanced across control, saline, and DCZ sessions. DCZ or saline was administered subcutaneously 1 h before the behavioral sessions. **e,** Color-coded spatial occupancy probability maps during the test phase for a representative saline-injected session (left) and a DCZ-injected session (right). White outlines in the arena denote the locations of the relocated objects within the arena. **f,** Bar plot showing the fraction of exploration time directed toward the relocated object during the test phase. Object preference was quantified across successive segments of cumulative object exploration time. Blue and red bars indicate saline- and DCZ-injected sessions, respectively. DCZ administration significantly reduced preference for the relocated object over the full exploration period ($p = 0.004$; Wilcoxon rank-sum test).

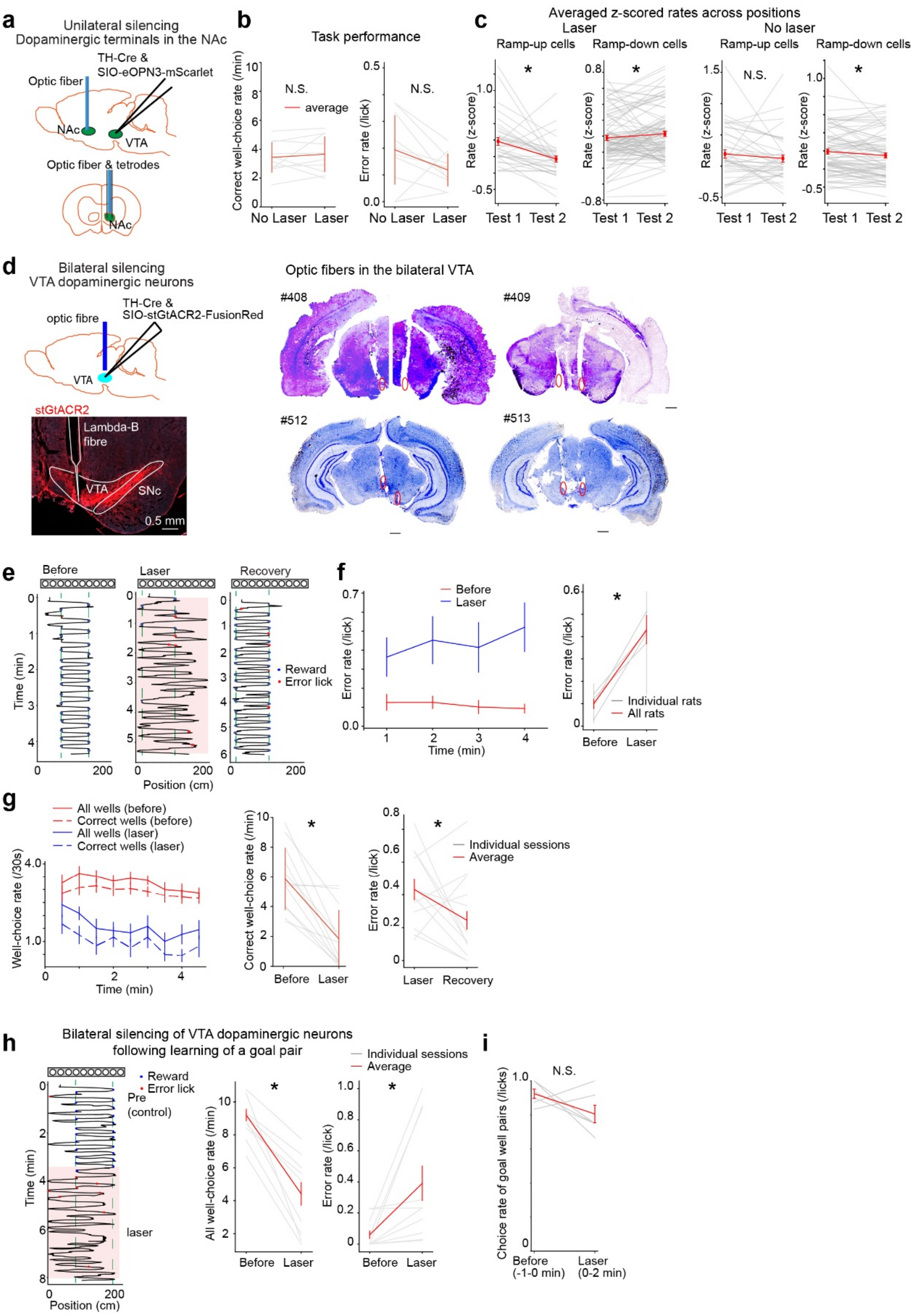


**Extended Data Figure 5. Impact of dopaminergic silencing on navigation performance.**
**a,** Schematic illustrating the strategy for unilateral silencing of dopaminergic inputs from the ventral tegmental area (VTA) to the nucleus accumbens (NAc). The axon-terminal–optimized inhibitory opsin eOPN3 was expressed in VTA dopaminergic neurons under the control of the tyrosine hydroxylase promoter. An AAV was injected unilaterally into the VTA, and a tetrode

microdrive combined with an optic fiber was implanted in the NAc of the same hemisphere to selectively inhibit dopaminergic terminals while recording NAc neuronal activity. **b,** Plots showing task performance, quantified as correct well-choice rate (left) and error rate (right). No significant differences were observed with laser application. **c,** Assessment of goal-directed firing in NAc neurons. Left: Comparison of goal-directed modulation during laser sessions, showing significant changes in both ramp-up and ramp-down populations (ramp-up; $n = 30$, ramp-down; $n = 64$, Wilcoxon signed-rank test, $p < 0.05$). Right: Comparison during no-laser sessions, showing no significant change in ramp-up cells ($n = 27$, Wilcoxon signed-rank test, $p = 0.336$) but a significant shift in ramp-down cells ($n = 41$, Wilcoxon signed-rank test, $p < 0.05$). **d,** Histological verification of viral expression and optic fiber placement in the VTA. Top left: Schematic illustrating the viral injection and fiber implantation strategy. The inhibitory opsin stGtACR2 was expressed in VTA dopaminergic neurons under the control of the tyrosine hydroxylase promoter. Bottom left: Representative coronal section showing stGtACR2 expression (red) and the optic fiber track in the VTA. Scale bar, 0.5 mm. Right: Nissl-stained coronal sections showing the locations of implanted optic fibers in the bilateral VTA. Red circles indicate fiber tip positions. Scale bar, 1 mm. **e,** Example behavioral trajectories showing the impact of VTA dopaminergic neuron silencing on navigation performance. Representative trajectories showing the animal's position over time before, during, and after laser application. Green dashed lines indicate goal locations; blue and red dots denote correct and error lick events, respectively; pink shaded regions indicate periods of laser illumination. **f,** Error rate calculated in 1 min moving windows for before (red) and laser (blue) sessions (12 sessions from 4 animals). Right: Summary of overall error rates across animals, showing a significant increase during laser application ($p < 0.05$; Wilcoxon rank-sum test). **g,** Plots showing well-choice performance across sessions. Left: Well-choice frequency calculated in 30 s moving windows, showing for all wells (solid lines) and correct wells (dashed lines) during before (red) and laser (blue) sessions (12 sessions from 4 animals). Middle: Comparison of correct well-choice rates between conditions; gray lines indicate individual sessions and red lines indicate session means. Right: Comparison of error rates between before and laser sessions. All comparisons were performed using the Wilcoxon rank-sum test (*$p < 0.05$). **h,** Impact of dopaminergic neuron silencing within the same goal-pair block. Left: Example trajectories showing animal navigation before and during laser application under an identical correct goal pair. Middle: Comparison of all well-choice rates between before and laser epochs (12 sessions from 2 animals). Right: Comparison of error rates between before and laser epochs, showing a significant increase during laser application ($p < 0.05$; Wilcoxon rank-sum test). **i**, Plot shows the discrimination performance between goal well pairs and other wells after silencing was applied following learning of the current goal pair. Unlike the error rate calculation in **h**, discrimination was quantified as the proportion of correct choices between goal pairs and other wells, irrespective of alternative choice sequences. Performance is shown before laser onset

(−1 to 0 min) and during laser stimulation (0 to 2 min) across daily sessions. Thin gray lines represent individual sessions, and red lines denote the mean ± s.e.m. No significant difference was observed ($p = 0.156$, Wilcoxon signed-rank test), indicating that post-learning silencing of VTA neurons did not significantly impair discrimination of the current goal pair from other wells.

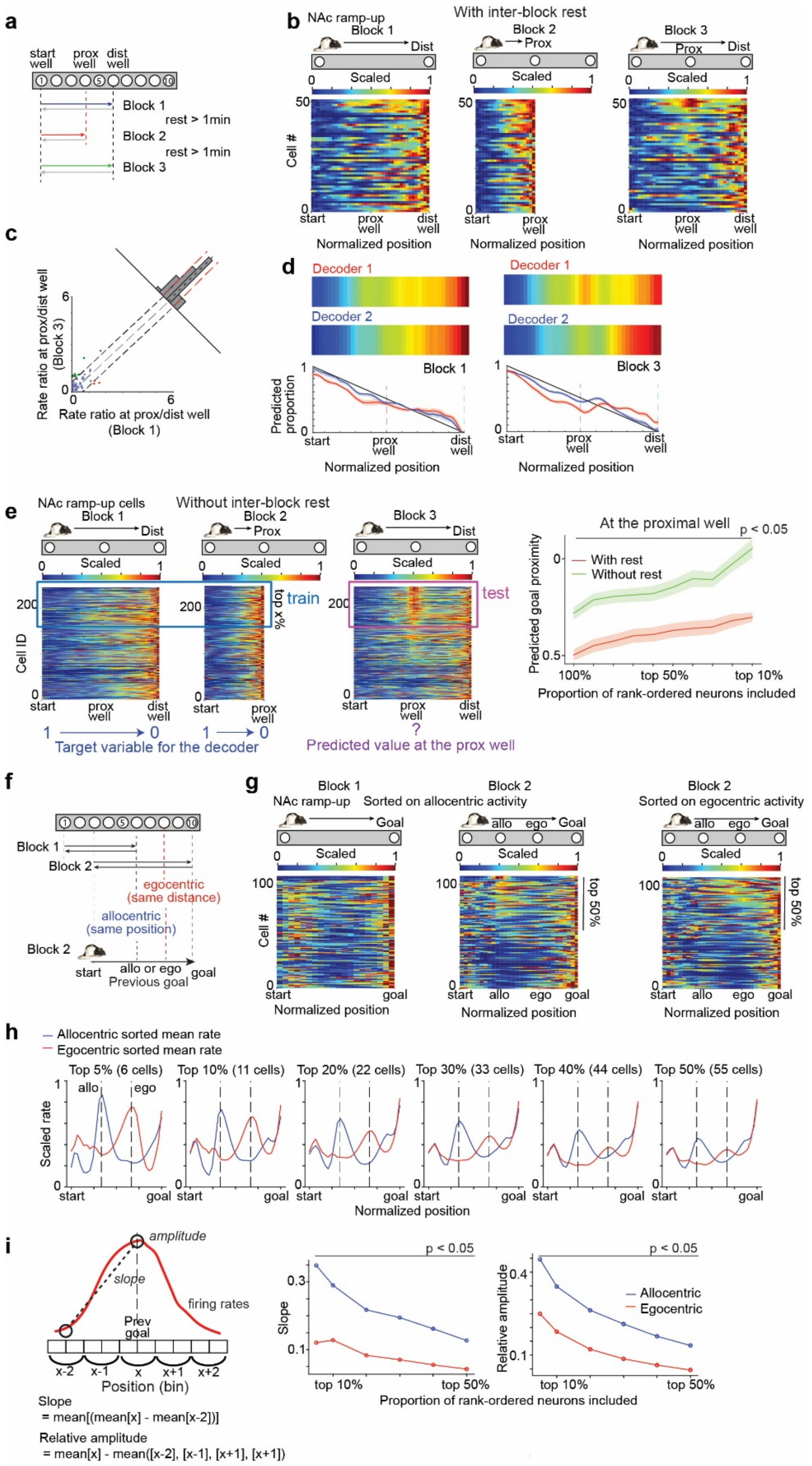


**Extended Data Figure 6. Latent coding of previous goals in the NAc reflects allocentric location and depends on task engagement.**

**a,** Schematic of the task design used to test the impact of an inter-block rest period on previous-goal coding in the NAc. Unlike the experiments in Fig. 4, animals were removed from the maze after each block and placed on a pedestal for at least 1 min before the next block. **b,** Color-coded firing-rate maps of NAc neurons exhibiting goal-directed ramp-up activity across blocks with an inter-block rest. Note the reduced firing at the proximal well in Block 3 compared with Fig. 4c. **c,** Difference in firing-rate ratios at the proximal versus distal wells between Blocks 1 and 3. **d,** Performance of proximity decoding for both current and previous goal location. As in Fig. 4d, two decoders were trained to decode either the previous goal (Decoder 1) or the current goal (Decoder 2). However, in Block 3, both decoders showed similar performance, in contrast to the results in Fig. 4e. **e,** Left: Schematic of the decoding procedure. Neurons were rank-ordered based on their activity at the proximal well location during Block 3, and subsets corresponding to different top-percentile ranges were selected for analysis. The color-coded maps shown are the same as Fig. 4c without the rest condition. A SVR decoder was trained to predict the proportional goal distance using neural activity from Blocks 1 and 2 and subsequently tested on neural activity from Block 3. Unlike the decoding procedure in Fig. 4d, the data from Block 3 were not used for decoder training. Right: Predicted goal proximity at the proximal well in Block 3 as a function of the proportion of neurons included in the decoder. For each condition, the decoder output was averaged across trials. Green indicates the without-rest condition, and red indicates the with-rest condition. Shaded areas represent the s.e.m. across trials. The black line at the top denotes proportions at which predicted proximity differed significantly between conditions ($p < 0.05$; Wilcoxon rank-sum test with Benjamini–Hochberg correction for multiple comparisons across bins). These results suggest that coding of the previous goal is significantly reduced following inter-block rest. **f,** Schematic of task design. Goal pairs for each block were assigned such that one goal from Block 1 was located between the goal pair used in Block 2. For Block 2 trials, two reference locations were defined based on Block 1 goals. The allocentric previous goal was defined as the absolute spatial location of the Block 1 goal, whereas the egocentric previous goal was defined as a distance-matched location obtained by translating the same start–goal distance from the new start position, dissociating allocentric and egocentric representations. **g,** Color-coded firing-rate maps showing the activity of individual NAc neurons during the journey. For Block 2 trials, neurons were sorted separately based on their activity at either the allocentric previous-goal location or the egocentric distance-matched location, while the data were identical. **h,** Rank-ordered average firing-rate profiles of NAc neurons sorted according to allocentric or egocentric previous-goal coding. Mean firing rates were computed over rank-ordered proportions of neurons. Allocentric-sorted averages (blue) consistently exhibited a sharper and more prominent peak than egocentric-sorted averages (red) across a wide range of selection thresholds. **i,** Quantification of the strength of previous goal coding in allocentric and egocentric coordinates. Two comparative metrics were computed for each neuron: the ramp-up slope to the target goal well and the relative peak amplitude at a target well compared to the surrounding spatial bins. The target well location (previous goal) was defined based on either allocentric or egocentric coordinates. The black line at the top denotes proportions at which predicted proximity differed significantly between conditions ($p < 0.05$; Wilcoxon rank-sum test with Benjamini–Hochberg correction for multiple comparisons across bins).

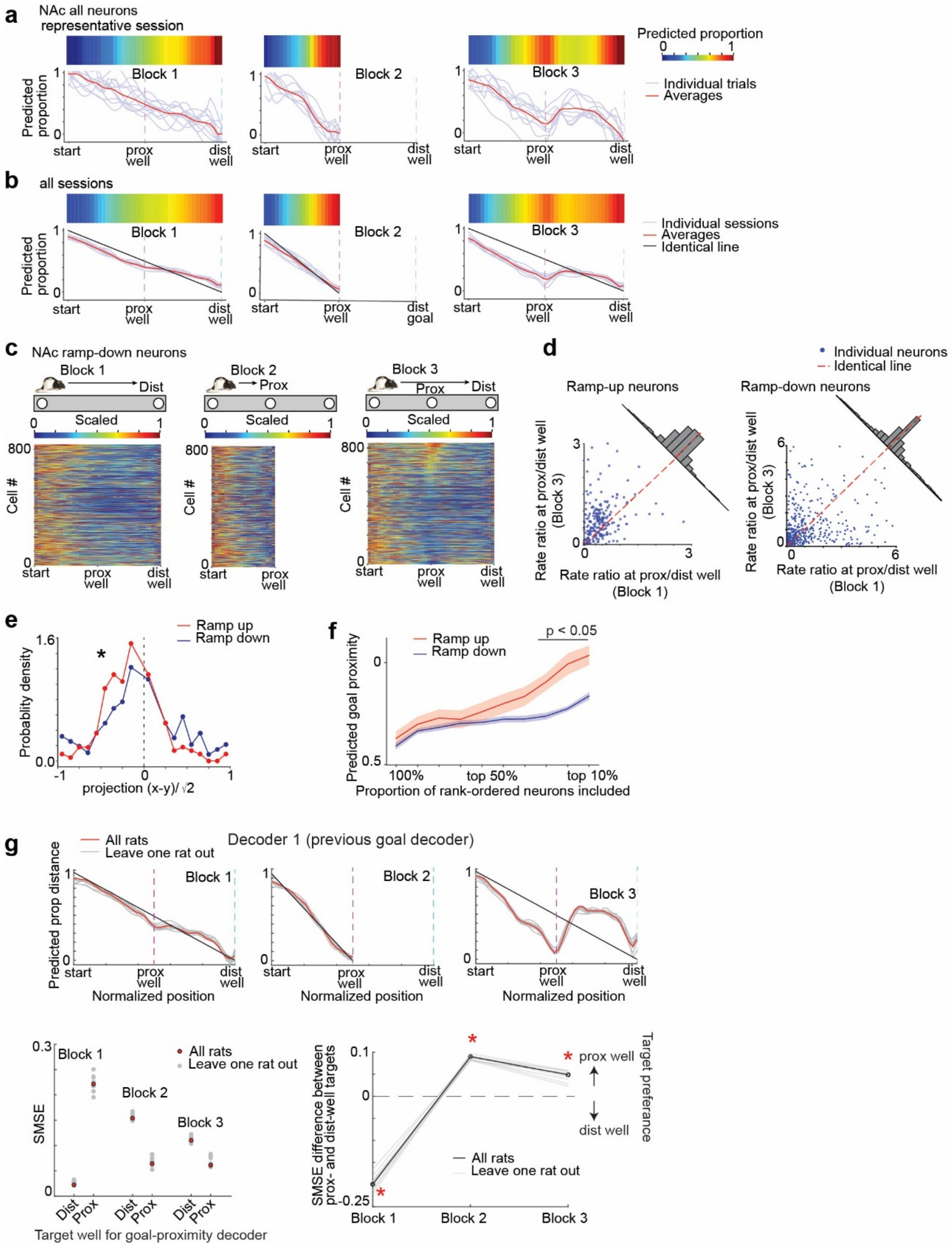


**Extended Data Figure 7. Previous-goal coding in the NAc is consistent across animals and structured by cluster-specific neuronal contributions.**

**a,** Example daily-session analysis showing previous-goal representations during outbound runs. The decoder was constructed using the same training and testing procedures as "Decoder 1" shown in Fig. 4d. This analysis, however, included all recorded neurons across clusters, not just ramp-up neurons. Predicted proportional distance decoded from neural population activity during outbound runs is shown as a color-coded plot (top) and as line plots (bottom), with individual trial-wise predictions shown in blue and their average shown in red. Results are shown separately for Blocks 1–3. **b**, Same as **a**, but showing session–averaged performance. Identical lines are shown in black. **c,** Color-coded plots showing normalized firing rates of individual goal-directed ramp-down neurons across blocks, corresponding to the top three panels of Fig. 4c. Neurons were sorted by activity at the previous well in Block 3. **d,** Scatter plots showing ratios of firing rates at the proximal versus distal well locations in Blocks 1 and 3 for ramp-up (left) or ramp-down (right) neurons. Data points are projected onto the diagonal axis to quantify relative changes in activity between the two well locations. **e,** Histograms of the values projected on the diagonal axis in **d** for ramp-up and ramp-down neurons. The distributions differed significantly between the two populations ($p < 0.001$; Kolmogorov–Smirnov test). **f,** Comparison of previous-goal representation strength between ramp-up and ramp-down neurons in the NAc. Decoder construction followed the schematic shown in Extended Data Fig. 6e (left). Predicted animal location at the proximal well in Block 3 is plotted as a function of the proportion of rank-ordered neurons included in the decoder. Blue indicates ramp-down neurons and red indicates ramp-up neurons. Shaded areas represent s.e.m. across trials. The black line at the top denotes proportions at which predicted proximity differed significantly between populations ($p < 0.05$; Wilcoxon rank-sum test with Benjamini–Hochberg correction for multiple comparisons across bins). These results indicate that previous-goal coding is significantly weaker in ramp-down NAc neurons compared to ramp-up neurons. **g,** Robustness of previous goal decoding across animals assessed using a leave-one-rat-out strategy. Decoding analyses were repeated while excluding data from one rat at a time. For each iteration, a decoder was trained to predict the previous goal in Block 3, using the same procedure as Decoder 1 described in Fig. 4d. Top: Decoder output aligned to normalized position for each block. Red traces show the averaged predictions using data from all animals, gray traces represent individual leave-one-rat-out iterations. Identical lines are shown in black. Leave-one-rat-out traces largely overlap with the all-rat results, indicating no systematic bias in decoding performance across animals. Bottom left: Decoding performance quantified as scaled mean squared error (SMSE) as in Fig. 4e. For each block, decoding error was evaluated relative to the proportional distance to either the proximal or distal well. Gray dots indicate leave-one-rat-out results, and red dots indicate the all-animal result. Bottom right: Target-well preference quantified as the differences in SMSEs between predictions of goal proximity to the proximal and distal wells. Black markers show the all-rat results, and gray lines show leave-one-rat-out results. Target-well preference was consistent across datasets, being below zero in Block 1 and above zero in Block 2 and 3 ($p = 0.002, 0.002, 0.002$ for Block 1-3, respectively; Wilcoxon signed-rank test).

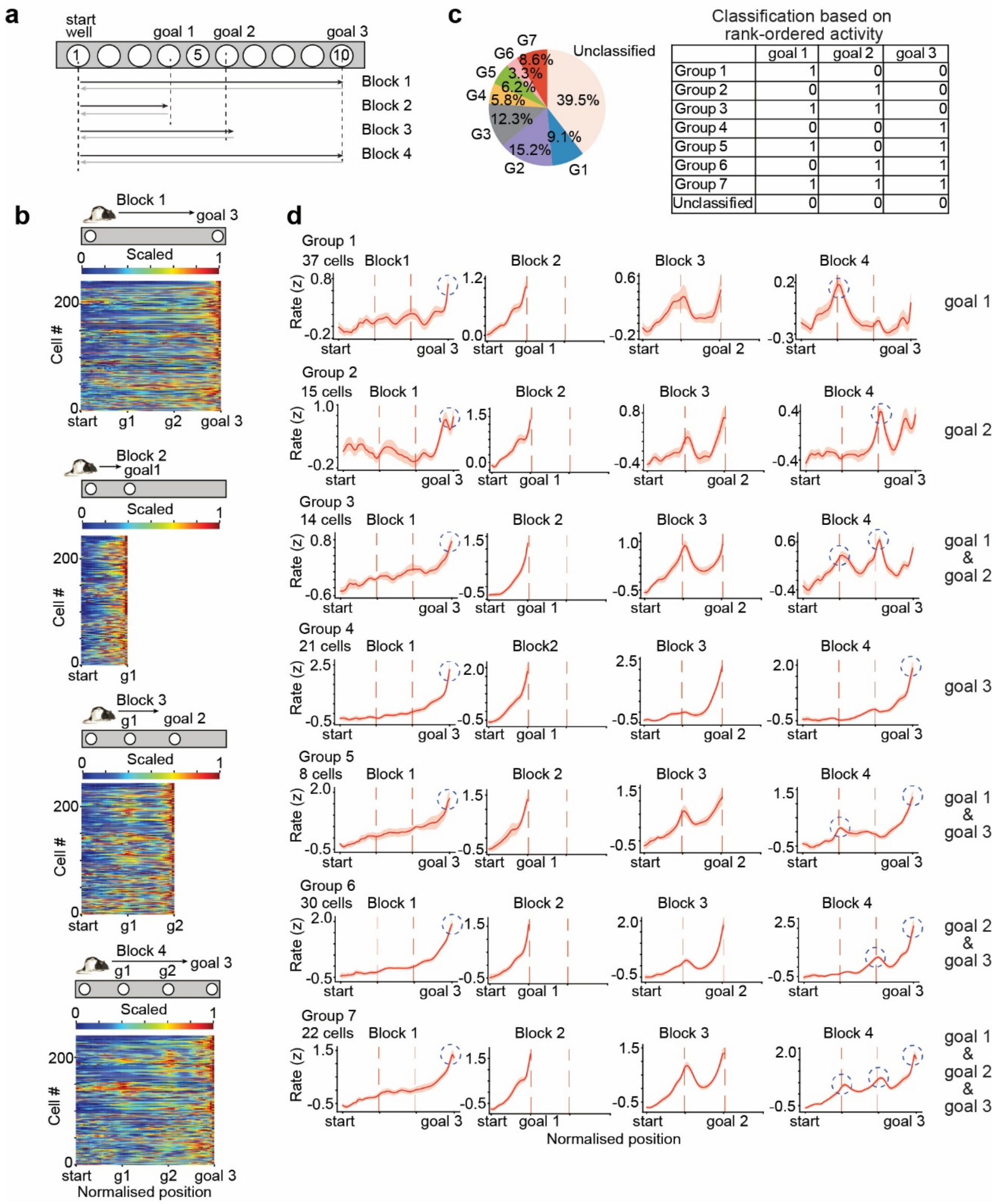


**Extended Data Figure 8. Previous goal coding in the NAc extends to multiple rewarded locations.**

**a,** Schematic of the task design to examine the coding of multiple previous goals. While the animals alternately navigate between a given pair of goals, the analysis focused on outbound runs. Across blocks, the animals navigated from a shared start well at the track end to one of three goal well locations (goal 1–3). In both Blocks 1 and 4, the animal navigated from the same start well to goal 3, but these blocks differed in the intervening experience of Blocks 2 and 3, targeting goal 1 and goal 2, respectively. Data from 6 rats, 28 daily sessions. **b,** Color-coded rate maps showing normalized firing rates of individual goal-directed ramp-up neurons

(243 cells) across task blocks. From top to bottom, colormaps correspond to Blocks 1–4. Neural activity was computed along normalized positions divided into three regions (start to goal 1, goal 1 to goal 2, and goal 2 to goal 3; 20 bins each). Neurons were classified based on activity during Block 4. For each neuron, mean firing rates were computed within each of the three regions. Neurons were ranked separately for each region by mean firing rate, and the top one-third of neurons for each region were selected. Neurons were then assigned to groups according to membership in these "top one-third" sets ($2^3$ = 8 possible patterns), with neurons belonging to none of the sets labeled unclassified. Neurons are ordered according to this grouping for visualization. **c,** Pie chart summarizing the proportion of ramp-up neurons assigned to each group. The schematic on the right summarizes the grouping strategy based on rank-ordered activity during Block 4. **d,** Mean firing rate profiles (mean ± s.e.m.) of ramp-up neurons for each group across task blocks, aligned to normalized position. The number of neurons contributing to each group is indicated. Blue-dashed circles in Blocks 1 and 4 highlight prominent activity peaks that serve as the basis for neuronal grouping. These peaks indicate that NAc neurons encode not only the currently targeted goal (goal 3) but also previously visited goals (goals 1 and 2), with individual neurons exhibiting distinct goal-specific biases. This pattern suggests that the decay of previous-goal representations cannot be explained solely by elapsed time. Rather, it reflects differential contributions of individual NAc neurons that maintain memory traces at specific goal locations, forming goal-location–specific representations with varying strengths and durations.

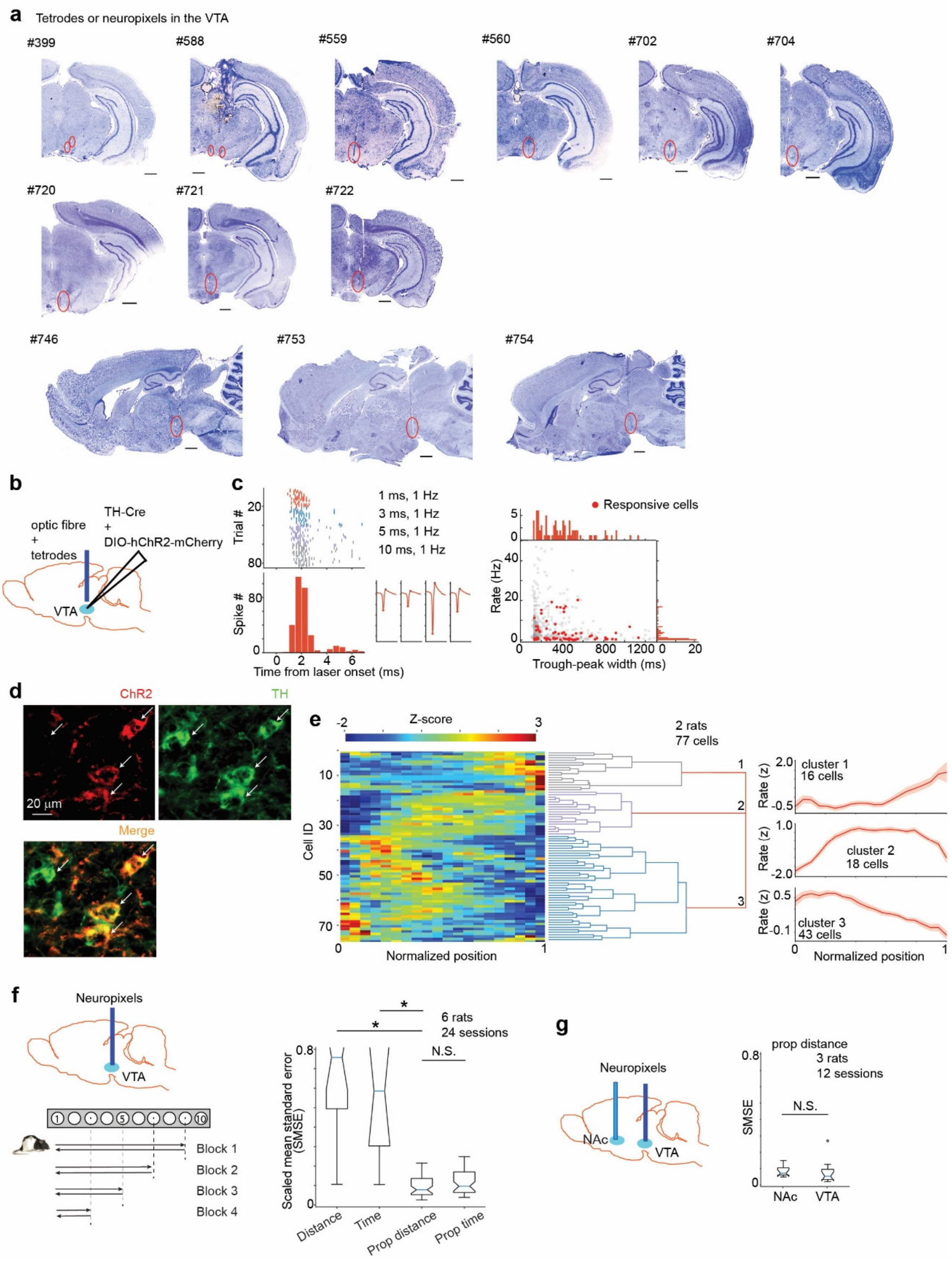


**Extended Data Figure 9. Optogenetic identification and functional characterization of VTA dopaminergic neurons**

**a,** Nissl-stained sections showing electrode implantation sites in the VTA. Coronal sections from 9 rats and sagittal sections from 3 rats are shown. Red circles indicate the locations of electrode tracks. Scale bar, 1 mm. **b,** Schematic illustration of the strategy for optogenetic tagging and recording from the same VTA dopaminergic (DA) neurons. An excitatory opsin

ChR2 was expressed in DA neurons under the control of Cre recombinase driven by the tyrosine-hydroxylase (TH) promoter. **c,** Example activity of an optogenetically identified DA neuron. Top, raster plots aligned to laser stimulation under different stimulation conditions (laser pulse duration: 1 ms, red; 3 ms, blue; 5 ms, purple; 10 ms, gray at 1 Hz; approximately 20 trials per condition). Bottom left, latency histogram pooled across all stimulation conditions. Bottom middle, spike waveforms recorded from a tetrode, showing four channels corresponding to a single isolated neuron. Bottom right, scatter plot of firing rate versus trough-to-peak spike width. Red dots indicate laser-responsive putative dopaminergic neurons, and gray dots indicate non-responsive neurons. Data from 2 rats, 77 laser-responsive cells. **d,** Immunohistochemical verification of ChR2 expression in VTA DA neurons. Top left: Fluorescence image showing ChR2 expression (red). Top right: Immunohistochemical staining for tyrosine hydroxylase (TH; green). Bottom: Merged image of two channels showing colocalization. White arrows indicate cells coexpressing ChR2 and TH. Scale bar, 20 μm. **e,** Hierarchical clustering of putative DA neurons based on their activity patterns during linear maze navigation. Left: Color-coded map showing z-scored firing rates of individual putative DA neurons (rows, cells; columns, normalized position from start to goal). Middle: Dendrogram illustrating clustering relationships among neurons. Right: Mean activity profiles for the three identified clusters. **f,** Top left: Schematic of Neuropixels probe implantation in the VTA. Bottom left: Task schematic (same as Fig. 1a). Right: Box plots showing SMSE of predicted proportional distance to the goal based on SVR. Four proximity variables—distance, time, proportional distance, and proportional time—were tested and compared. SMSEs for proportional distance were significantly lower than distance ($p < 0.001$; Wilcoxon rank-sum test) and time ($p < 0.001$) but did not differ significantly from proportional time ($p = 0.081$). Data from 6 rats, 24 daily sessions. **g,** Left: Schematic of simultaneous Neuropixels recordings from the NAc and VTA. Right: Box plots comparing SMSEs for proportional distance decoding between NAc and VTA population activity. No significant difference was observed ($p = 0.225$; Wilcoxon rank-sum test). Data from 3 rats, 12 daily sessions.

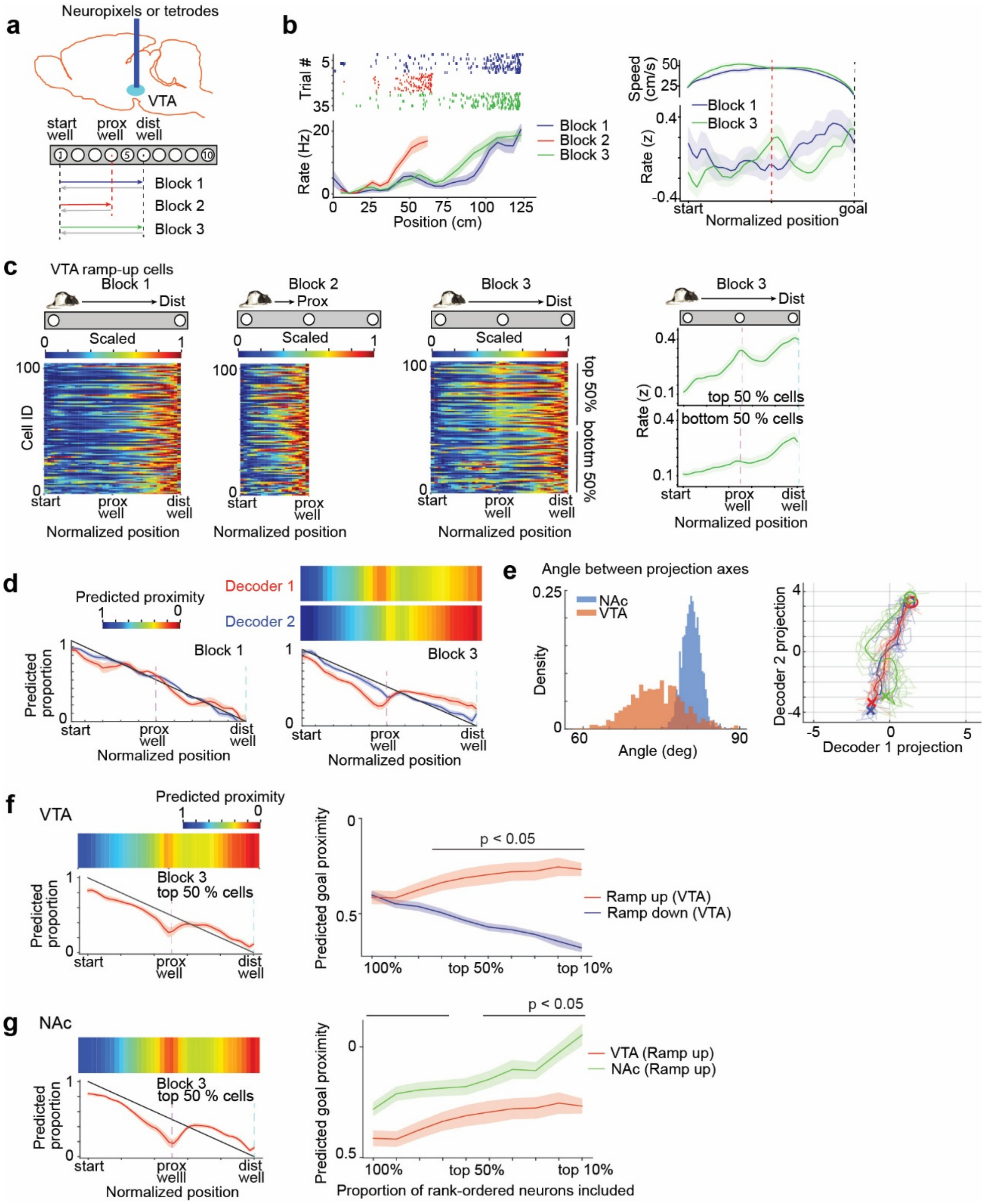


**Extended Data Figure 10. Limited evidence for previous-goal coding in the VTA**

**a,** Top, schematic illustration of Neuropixels probe implantation in the VTA. Bottom, task schematic (same as Fig. 4a). Data from 9 rats, 25 daily sessions. **b,** Same as Fig. 4b, but shown for a representative VTA neuron. Left: Firing rates across three blocks during outbound runs. Right: Average firing rates of VTA neurons exhibiting goal-directed ramp-up firing and top 25% previous-goal well activity, computed under matched running speeds between Blocks 1 and 3. Trials were iteratively excluded to match running speed distributions between blocks, and population activity was obtained by averaging firing rates across neurons during the matched trials. Plots show running speed (top) and firing rate (bottom) for Blocks 1 and 3.

Positions were normalized; the red dashed line indicates the proximal well location, and the black dashed line indicates the distal well location. **c,** Color-coded rate maps showing normalized firing rates of individual VTA neurons across blocks, as in Fig. 4c. Neurons were sorted by their activity at the proximal well in Block 3. Right: Mean firing rates across neurons in the top 50% or bottom 50% based on the activity at the proximal well in Block 3. **d,** Construction of decoders for estimating proportional distance to the previous goal (Decoder 1) or the current goal (Decoder 2) based on VTA population activity, as in Fig. 4d,e. Predicted proportional distance to the previous goal during outbound runs is shown for Blocks 1 and 3. Note the weak proximity prediction to the previous goal by Decoder 1, in contrast to the corresponding result from the NAc in Fig. 4e. **e,** Orthogonality of decoding axes. Left: Comparison of angle distributions between projection vectors of Decoder 1 (proportional distance to the previous goal) and Decoder 2 (proportional distance to the current goal) derived from NAc (blue) and VTA (red) activity. Distributions were obtained using permutation analysis and differed significantly between regions ($p < 0.001$; Kolmogorov–Smirnov test), indicating that the two decoders constructed from VTA activity are less orthogonal. Right: Population VTA activity dynamics during navigation plotted onto the two decoder axes. Note that the separation of trajectories in Block 3 relative to Blocks 1 and 2 is less pronounced than in the corresponding NAc results shown in Fig. 4f. **f,** Comparison of previous-goal representation strength between ramp-up and ramp-down neurons in the VTA. Left: Decoding of proportional goal distance using the top 50% of rank-ordered VTA neurons in **c** during outbound runs in Block 3 (top, color-coded plots; bottom, mean ± s.e.m.). Decoder construction followed the schematic shown in Extended Data Fig. 6e (left) using Blocks 1 and 2 for training and Block 3 for testing. Right: Same as in Extended Data Fig. 7f, but for VTA neurons. Blue indicates ramp-down neurons and red indicates ramp-up neurons. Shaded areas represent s.e.m. across trials. The black line at the top denotes proportions at which predicted proximity differed significantly between populations ($p < 0.05$; Wilcoxon rank-sum test with Benjamini–Hochberg correction for multiple comparisons across bins). **g,** Comparison of previous-goal coding strength between ramp-up neurons in the NAc and VTA. Left: Decoding performance using the top 50% of rank-ordered NAc ramp-up neurons based on previous goal activity (same procedure as in **f**). Right: Predicted goal proximity at the proximal well as a function of the proportion of rank-ordered neurons included, comparing VTA (red) and NAc (green). Axes, shading, and statistical annotations are the same as in **f**. Together with photometry measurements of dopamine concentration in the NAc (Fig. 4g,h), these results indicate that previous-goal coding is substantially weaker in the VTA than in the NAc.
